# Case-only exome variation analysis of severe alcohol dependence using a multivariate hierarchical gene clustering approach

**DOI:** 10.1101/2022.03.16.484608

**Authors:** Amanda E. Gentry, Jeffry C. Alexander, Mohammad Ahangari, Roseann E. Peterson, VCU Alcohol Research Center working group, Michael F. Miles, Jill C. Bettinger, Andrew G. Davies, Mike Groteweil, Silviu A. Bacanu, Kenneth S. Kendler, Brien P. Riley, Bradley T. Webb

## Abstract

**Background:** Variation in genes involved in ethanol metabolism has been shown to influence risk for alcohol dependence (AD) including protective loss of function alleles in ethanol metabolizing genes. We therefore hypothesized that people with severe AD would exhibit different patterns of rare functional variation in genes with strong prior evidence for influencing ethanol metabolism and response when compared to genes not meeting these criteria.

**Objective:** Leverage a novel case only design and Whole Exome Sequencing (WES) of severe AD cases from the island of Ireland to quantify differences in functional variation between genes associated with ethanol metabolism and/or response and their matched control genes.

**Methods:** First, three sets of ethanol related genes were identified including those a) involved in alcohol metabolism in humans b) showing altered expression in mouse brain after alcohol exposure, and altering ethanol behavioral responses in invertebrate models. These genes of interest (GOI) sets were matched to control gene sets using multivariate hierarchical clustering of gene-level summary features from gnomAD. Using WES data from 190 individuals with severe AD, GOI were compared to matched control genes using logistic regression to detect aggregate differences in abundance of loss of function, missense, and synonymous variants, respectively.

**Results:** Three non-independent sets of 10, 117, and 359 genes were queried against control gene sets of 139, 1522, and 3360 matched genes, respectively. Significant differences were not detected in the number of functional variants in the primary set of ethanol-metabolizing genes. In both the mouse expression and invertebrate sets, we observed an increased number of synonymous variants in GOI over matched control genes. Post-hoc simulations showed the estimated effects sizes observed are unlikely to be under-estimated.

**Conclusion:** The proposed method demonstrates a computationally viable and statistically appropriate approach for genetic analysis of case-only data for hypothesized gene sets supported by empirical evidence.

## Introduction

Alcohol use disorder (AUD) is a common, moderately heritable disorder with significant social and economic impact. Twin(1–11), family (12), and adoption studies(13–16) consistently show that genetic influences have a large impact on the risk for AUD and alcohol-related phenotypes, with twin-based heritability estimates of ~0.50(17) and SNP-based heritability estimates of ~0.056(18). In recent years, genome-wide association studies (GWAS) have successfully identified common single nucleotide variants (cSNV) robustly associated with alcohol consumption(19), the Alcohol Use Disorders Identification Test (AUDIT)(20), and problematic alcohol use(18,21). Many of these identified cSNVs impact genes encoding the alcohol metabolizing enzymes such as the cluster of alcohol dehydrogenase (*ADH*) genes on chromosome 4. Additionally, there is accumulated evidence that variation in *CYP, CAT*, and *ALDH* genes are also involved in alcohol metabolism and AUD risk. *CYP2E1* expression is induced by chronic alcohol consumption and is thought to contribute to ethanol metabolism in the brain where *ADH* activity is limited(22). *CAT* is also widely expressed in the brain and a number of studies suggest that polymorphisms in the *CAT* gene are involved in the level of response to ethanol and alcohol dependence (AD) and abuse(23,24). *CYP2E1* and *CAT* are considered part of the canonical set of genes contributing to oxidative metabolism of ethanol(25).

While the effect of variants on *ADH, CYP2E1*, and *CAT* genes are more subtle, the well documented *ALDH2*2* (rs671) loss of function (LOF) allele shows only 20-40% of wild type enzymatic activity in heterozygote carriers due to the homo-tetrameric structure of mature *ALDH2*. This variant is common in individuals of East Asian ancestry, but largely absent in other populations(26) and is associated with lower rates of alcohol abuse/dependence(27) because the reduced enzymatic activity leads to accumulation of acetaldehyde and unpleasant symptoms such as excessive flushing in carriers.

Although significant progress has been made in cSNV identification in AUD and alcohol-related phenotypes, rare or intermediate frequency single nucleotide variant (rSNV) investigations are largely limited by sample power. Findings from the 1000 Genomes Project suggest that rare functional variation is frequent in the genome with approximately 400 premature stop, splice-site disrupting and frame-shift alleles affecting 250-300 genes per individual genome(28) and display strong population specificity(29). Sequencing studies in psychiatric disorders suggest that rare functional variation is an important element of risk for intellectual disability(30), autism spectrum disorders(31), and schizophrenia(32). While rare functional variants have not yet been widely studied through exome sequencing in AUD, early evidence points to their significant contribution to the genetic architecture of alcohol use phenotypes(18,20,33,34). Furthermore, studies across psychiatric phenotypes such as autism spectrum disorders(31), and schizophrenia(32) show an excess rate of rSNV in genes identified from common variant GWAS signals in cases compared to controls, suggesting that there is a convergence between cSNV and rSNV signals and disease risk is likely influenced by multiple alleles of varying frequencies in the same loci(35).

As a complement to genetic studies in humans, mice and invertebrate model organisms can also facilitate the identification of genetic mechanisms or orthologous genes that influence AUD in humans. Studies in mice identified genes showing altered expression in prefrontal cortex (PFC) after intraperitoneal injection of 1.8 g/kg of ethanol versus saline control, and identified a subset of these as hub genes defined by both high connectivity and high centrality in coexpression networks(36). Introduction of mutations or knockdown strategies in invertebrate models such as *D. melanogaster* or *C. elegans* can also identify genes involved in behavioral response to ethanol(37–39).

In this study, we sought to investigate exome variation in a sample of 190 severely affected alcohol dependence (AD) cases from the island of Ireland using a novel case-only analysis framework. Because of previous evidence of the protective effects of LOF alleles in ethanol metabolizing genes, we hypothesized that individuals with severe AD would show less functional coding variation in these genes compared to control genes with similar attributes. Furthermore, we extended this work to test sets of genes identified in model organisms and also hypothesized that genes robustly shown to impact alcohol-related outcomes in mice and invertebrates would show divergent patterns of exome variation in comparison to control genes with similar attributes. For each hypothesis, we sought to compare the numbers of LOF, missense (MIS), and synonymous (SYN) variants between genes of interest and a matched set of control genes. Given the absence of implemented methods to test our hypotheses in a case-only framework with related subjects, we sought to develop a novel framework to address these challenges. As an alternative to comparing aggregate exome variation between cases and controls, we matched genes of interest to control genes with similar attributes using gnomAD(40) as an external source of independent information, and unbiased comparisons across sets were made in a case-only analysis framework. Since comparison sets would be derived from within an individual’s genome, they would not be subject to any potential inflation from stratification due to population structure or other sources of bias. We present this framework as a complementary method to case-control association designs that can be performed in samples without matched controls and provides a rigorous framework for hypothesis testing where robust prior sources of evidence are available.

## Materials and Methods

### Sample description

The Irish Affected Sib-Pair Study of Alcohol Dependence (IASPSAD) sample(8) was collected from 1998-2002 in treatment facilities and hospitals in the Republic of Ireland and Northern Ireland. Probands with all four grandparents born in Ireland or Britain were ascertained for a diagnosis of DSM-IV AD(41) with one or more affected siblings. Lifetime history of AD was assessed using a modification of the Semi-Structured Assessment for the Genetics of Alcoholism (SSAGA) version 11(42) which permits evaluation of International Classification of Disease (ICD)-10, Feighner(43), RDC(44), DSM-III-R(45) and DSM-IV diagnostic criteria. The sample is severely affected, with ~87% of probands and ~78% of siblings endorsing ≥6 of the 7 DSM-IV AD criteria and 92% reported withdrawal symptoms. Parents were evaluated for lifetime history of alcohol abuse and AD based on the Structured Clinical Interview for DSM (SCID)(46), the CAGE Assessment (<u>C</u>utting Down, <u>A</u>nnoyance by Criticism, <u>G</u>uilty Feeling, and <u>E</u>ye Openers)(47), and Fast Alcohol Screening Test (FAST) items developed to screen for drinking problems(48).

### Exome capture and sequencing

Exome capture was performed using the Agilent SureSelect V5 71Mb exome + untranslated regions target kit, followed by library preparation and sequencing on the Illumina HiSeq X Ten system at BGI. The 190 samples were sequenced in 3 batches, with the first (n=57) and second (n=76) batches via 2×90 and the third batch (n=57) via 2×100 paired end sequencing.

### Variant calling

Sequence data was processed and called according to GATK3 best practices and summarized for quality control using FastQC (v 0.11.4). Sequence read alignment was performed using BWA-MEM (v 0.7.12) to hs37d5 reference genome, followed by reordering, duplicate marking and insertion deletion realignment with Picard (v 2.0.1). Variant calling was done using HaplotypeCaller, and variant quality score recalibration was carried out using Variant Quality Score Recalibration (VQSR) in GATK (v 3.5).

### Annotation

SnpEff (v 4.3, database GRCh37.75) was used to obtain gene annotations and included only the transcripts found in the gnomAD release 2.1.1 gene constraints.

### Description of candidate genes of interest

Selection of genes of interest (GOI) was carried out in collaboration with investigators from the Virginia Commonwealth University Alcohol Research Center (VCU-ARC), focused on cross-species discovery and functional interpretation of genes involved in AUD and related phenotypes. Three sets of alcohol related GOI were constructed for this analysis which crossed three taxonomic categories including human, mouse, and invertebrates. The first GOI set contains 11 ethanol metabolizing genes whose products are known to be involved with ethanol metabolism in humans. These included the *ADH* genes (n=7), *ALDH1A1, ALDH2, CAT*, and *CYP2E1* (Table 1). The second GOI set contains 109 hub genes with both high connectivity and high centrality in co-expression networks that show altered expression in mouse PFC 4 hours after intraperitoneal injection of 1.8 g/kg of ethanol versus saline control(36) (Supplemental Table S1). The third GOI set contains 358 genes for which manipulation in invertebrate model organisms results in altered ethanol response phenotypes(49) (Supplemental Table S2).

**Table 1:** Ethanol-metabolizing genes of interest (n=11), with annotation indicating whether they were present in the gnomAD gene constraints file (10), in the mouse brain expression set (1), and in the invertebrate set (5).

| Gene | Transcript | Present in gnomAD | Present in the invertebrate GOI set | Present in the mouse brain expression GOI set |
| --- | --- | --- | --- | --- |
| ADH1A | ENST00000209668 | TRUE | TRUE | FALSE |
| ADH1B | ENST00000305046 | TRUE | TRUE | FALSE |
| ADH1C |  | FALSE | TRUE | FALSE |
| ADH4 | ENST00000265512 | TRUE | FALSE | FALSE |
| ADH5 | ENST00000296412 | TRUE | FALSE | FALSE |
| ADH6 | ENST00000394899 | TRUE | FALSE | FALSE |
| ADH7 | ENST00000476959 | TRUE | FALSE | FALSE |
| ALDH1A1 | ENST00000297785 | TRUE | TRUE | FALSE |
| ALDH2 | ENST00000261733 | TRUE | TRUE | TRUE |
| CAT | ENST00000241052 | TRUE | FALSE | FALSE |
| CYP2E1 | ENST00000463117 | TRUE | FALSE | FALSE |

### Annotations for gene clustering

In order to create matched sets of genes for our case-only analytic approach, we utilized gene-level annotation information from the gnomAD database (v 2.11)(40). The seven gnomAD annotations used to cluster all genes included: (1–3) the ratios of observed to expected (O/E) counts for each variant class (LOF, MIS, and SYN), (4–6) a z-score for each O/E ratio, and (7) the probability of loss of function intolerance (pLI) score. In addition to these metrics from gnomAD, genes were annotated for clustering with metrics for (8) genomic length, (9) transcript length, and (10) number of exons. In total, ten annotations were utilized for clustering. While correlations between some variables were high (see Table 2), none were considered close enough to to warrant dropping from clustering.

**Table 2:** Correlations between the gene metrics from the gnomAD database.

|  | O/E<br>LOF | O/E<br>MIS | O/E<br>SYN | LOF<br>z-score | MIS<br>z-score | SYN<br>z-score | pLI | Gen.<br>Length | Tran.<br>Length | No. of<br>Exons |
| --- | --- | --- | --- | --- | --- | --- | --- | --- | --- | --- |
| O/E<br>LOF | 1 | 0.551 | 0.207 | -0.728 | -0.548 | -0.137 | -0.619 | -0.141 | -0.156 | -0.161 |
| O/E<br>MIS | 0.551 | 1 | 0.416 | -0.472 | -0.874 | -0.395 | -0.495 | -0.062 | 0.002 | -0.071 |
| O/E<br>SYN | 0.207 | 0.416 | 1 | -0.113 | -0.326 | -0.848 | -0.031 | -0.007 | 0.008 | -0.028 |
| LOF<br>z-score | -0.728 | -0.472 | -0.113 | 1 | 0.648 | 0.069 | 0.654 | 0.335 | 0.572 | 0.59 |
| MIS<br>z-score | -0.548 | -0.874 | -0.326 | 0.648 | 1 | 0.403 | 0.564 | 0.135 | 0.149 | 0.245 |
| SYN<br>z-score | -0.137 | -0.395 | -0.848 | 0.069 | 0.403 | 1 | 0.016 | -0.032 | -0.101 | -0.03 |
| pLI | -0.619 | -0.495 | -0.031 | 0.654 | 0.564 | 0.016 | 1 | 0.148 | 0.158 | 0.141 |
| Gen.<br>Length | -0.141 | -0.062 | -0.007 | 0.335 | 0.135 | -0.032 | 0.148 | 1 | 0.315 | 0.378 |
| Tran.<br>Length | -0.156 | 0.002 | 0.008 | 0.572 | 0.149 | -0.101 | 0.158 | 0.315 | 1 | 0.802 |
| No. of<br>Exons | -0.161 | -0.071 | -0.028 | 0.59 | 0.245 | -0.03 | 0.141 | 0.378 | 0.802 | 1 |

O/E: Observed/Expected
Gen. Length: Genomic Length
Tran. Length: Transcript Length
No. of Exons: Number of Exons

### Gene sets for analysis

Given that gnomAD metrics were needed to annotate the GOI, we had to drop 1 gene from the human ethanol metabolizing set because it did not appear in gnomAD (Table 1), 1 gene from the mouse hub gene set (Supplemental Table S1), and 5 genes from the ethanol behavioral response in invertebrates set (Supplemental Table S2). The final GOI sets contained 10, 108, and 353 genes in the human, mouse, and invertebrate sets, respectively, for inclusion in clustering and subsequent analyses. For hypothesis testing, we considered 3 testing sets: Set 1 contained only the 10 GOI for human ethanol metabolization, Set 2 contained Set 1, plus the addition of the 108 mouse hub genes. One gene appeared in both the human and mouse sets, therefore Set 2 contained a total of 117 genes. Set 3 contained Set 2, plus the addition of the ethanol behavior response in invertebrates set. One hundred and eleven of the invertebrate set genes appeared in Set 2, therefore Set 3 contained a total of 359 genes. Figure 1 shows the flow chart of filtering and sample sizes (panel a), as well as the Venn diagram (panel b) illustrating overlap between the three GOI sets, after removing genes absent from gnomAD.

**Figure 1:**
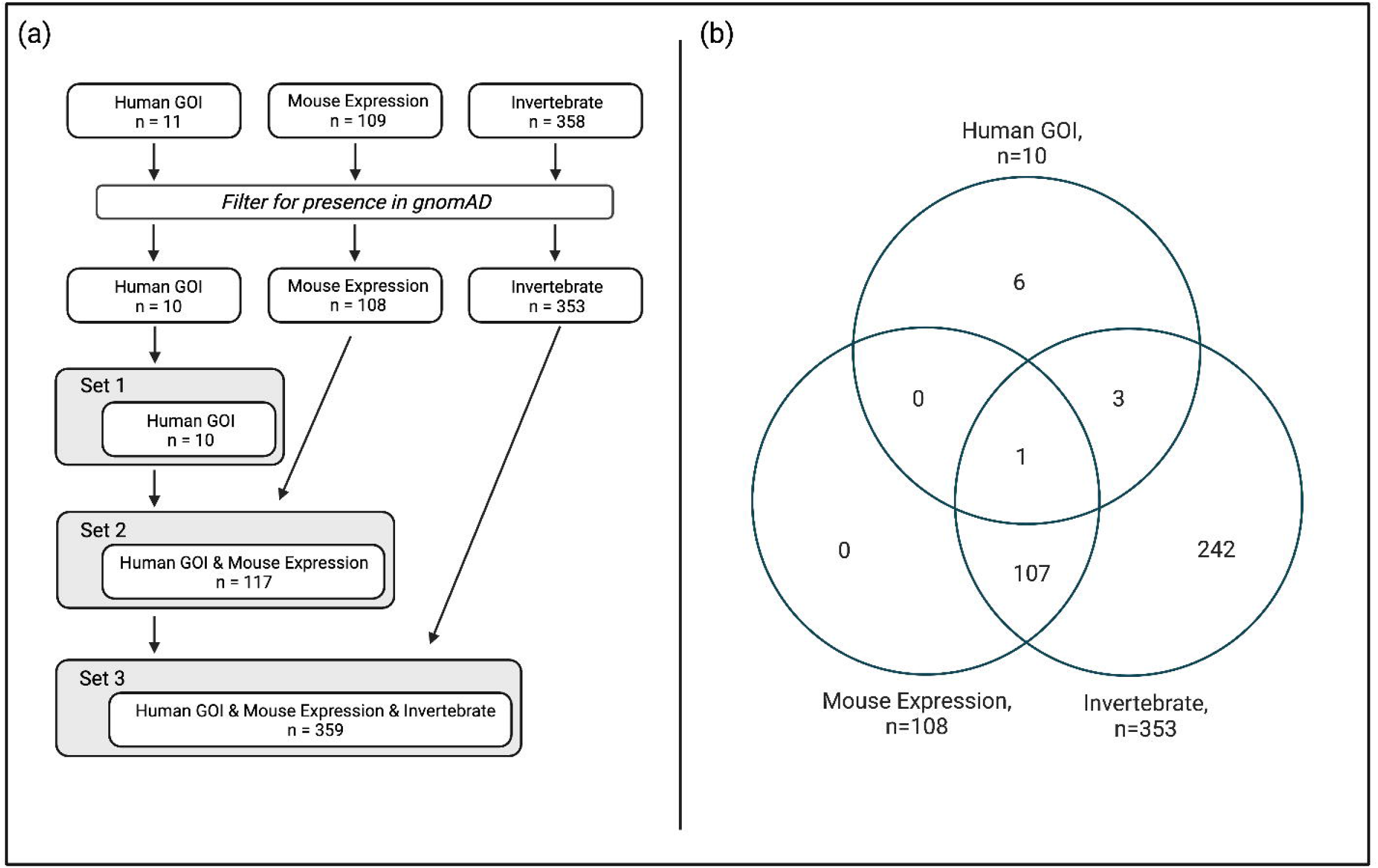
(a) Flow diagram with gene counts for the human GOI set, the mouse expression set, and the invertebrate set, through filtering and combination for hypothesis testing sets, and (b) Venn diagram illustrating the overlap of genes in the sets of interest, after filtering for presence in gnomAD. (*Figure created with BioRender.com*)

### Clustering

Multivariable single linkage agglomerative clustering of all canonical gnomAD genes was carried out using the *hclust* package (method =“single”) in R (v 3.6.0). The clustering algorithm was used to identify genes similar to each GOI based on the gene-level variables described above to create a control set of genes against which the GOI could be compared in a regression framework. This procedure utilizes information from gnomAD only, separate from the IASPSAD sample exomes. Clustering was performed on 10 gene metrics using a dissimilarity metric of 1 – |*cor*(*X*)|, where *X* is the *q* × *p* matrix of the *q* = 10 normalized metrics of each gene (*p* = 19,108). Figure 2 provides an example of gene clusters plotted according to three of these gnomAD gene metrics (SYN z-score, transcript length, and pLI). The data shown in this figure represent 4 gene clusters (shown in different colors) and a set of genes chosen at random (in gray), with ellipses highlighting the shape of the cluster. Matrix *cor(X*)represents the full set of pairwise correlations for all 19,108 genes in the set.

**Figure 2:**
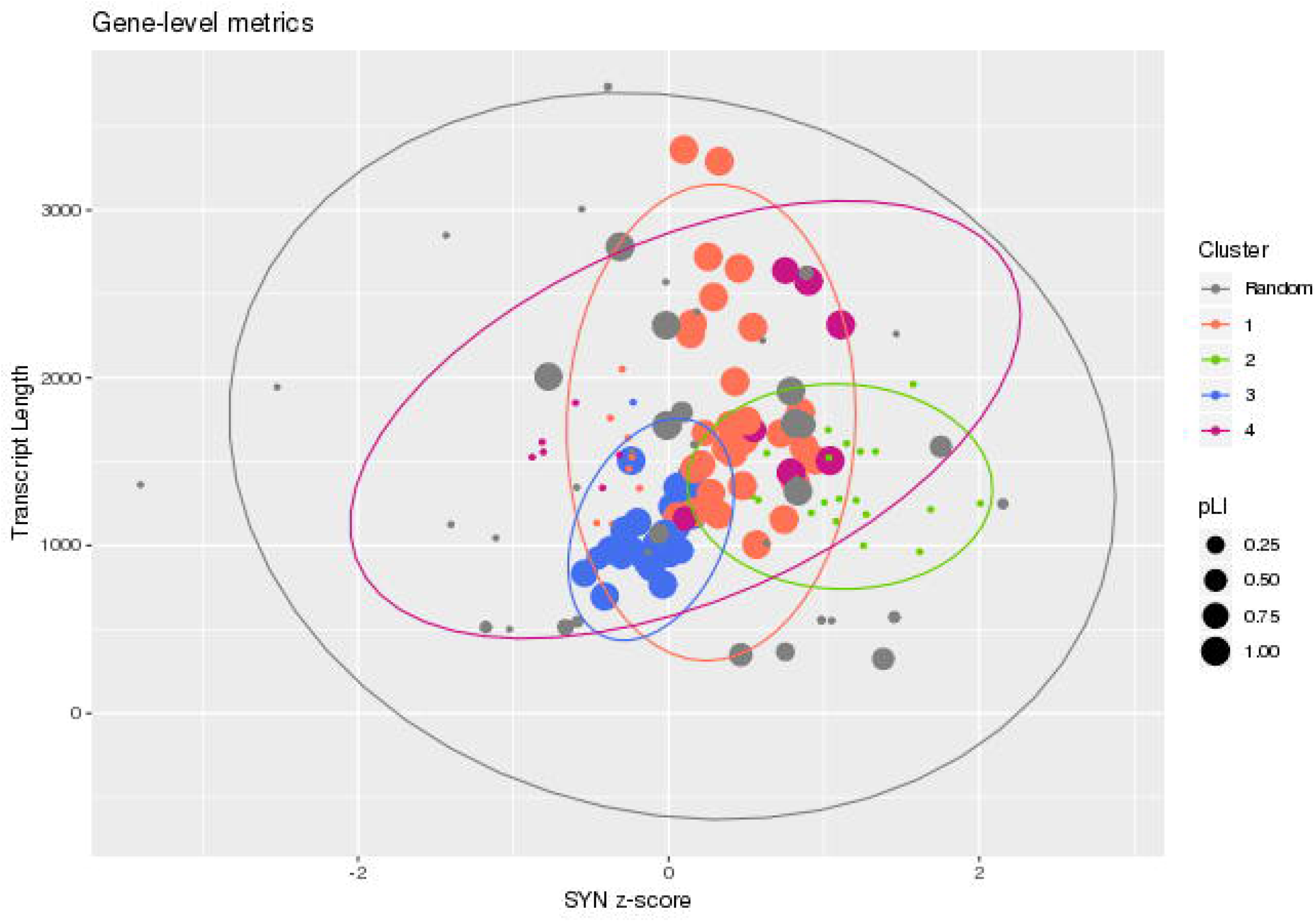
Example gene clusters plotted to show three of the 10 clustering gene metrics, SYN z-score, transcript length, and pLI. The 4 clusters in orange, green, blue, and pink represent actual gene clusters from the data, while the genes in gray represent a random selection of genes.

The hierarchical clustering was carried out in a stepwise fashion, beginning with all observations (genes) in a separate cluster of their own. At each successive step, the two least dissimilar clusters were joined together. We then pruned the final trees for all three GOI sets, choosing a height with enough genes clustered with each GOI for useful comparison while not exceeding 20% of the exome included in the final branches. Branches in the tree which contain at least one GOI were termed Clusters of Interest (COI). After a cut height is chosen, all other clusters not containing a GOI were pruned from the tree.

### Association testing

We used logistic regression to compare observed counts of LOF, MIS, and SYN variants aggregated from the IASPSAD sample exome data between GOI and the matched control genes. In this framework, instead of human subjects, the dependent variable is the gene which is either a GOI for a given hypothesis or not and coded 1 or 0, respectively. Therefore, the logistic regression quantifies the probability that a given gene is a GOI as a function of the observed counts of LOF, MIS, and SYN variants aggregated across the subjects in the IASPSAD sample. In other words, this framework tests whether or not the number of LOF, MIS, and/or SYN variants contributes to the probability that a given gene contributes to a hypothesized gene set such as alcohol metabolism. For each of the three GOI sets, the model is constructed as follows:

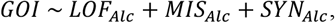

where GOI represents case/control status of the genes as described above for each set, and *LOF_Alc_*, *MIS_Alc_*and *SYN_Alc_* represent the observed counts of LOF, MIS, and SYN variants in that GOI, respectively. This model tests a null hypothesis of no association between the LOF, MIS, and SYN variant counts and probability of being a GOI versus a non-GOI. Evidence against the null hypothesis indicates that the alcohol-related genes of interest contain differing amounts of variation, as measured by counts of LOF, MIS, and SYN variants, from their matched control genes within the alcohol sample.

### Simulations

To empirically demonstrate the utility of this approach, we applied the method to a series of simulated datasets and assessed their performance. These simulations were conducted post-hoc and designed to follow distributional patterns and estimated effects observed in the real data model for the overarching purpose of increasing confidence in the real data results. Full details of the simulation parameters are given in the Supplemental Section S1 and Supplemental Table S3, but in brief, the simulations replace the observed LOF, MIS, and SYN variant counts for all 19,108 genes from the sample data with randomly generated data according to the following steps:

1. Set the distributions of the simulated data mimicking the observed distributions of LOF, MIS, and SYN variant counts in the alcohol sample data according to negative binomial distributions
2. Simulate the random LOF, MIS, and SYN data using the *SimCorrMix* package, according to the correlation structure in the observed alcohol sample data, such that:

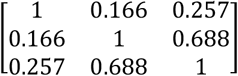
3. Generate the GOI probabilities using a logit model with beta effects for the intercept, LOF, MIS, and SYN values set at (−3, 0.75, 0.05, 0.1), (−5, 1.25, 0.1, 0.2), or (−7, 1.75, 0.2, 0.3). These values were chosen to broadly reflect the estimated effects observed in the data.
4. Generate random error according to a normal distribution with mean = 0 and standard deviation ranging from 0.55-1.45
5. Utilizing the clusters generated from gnomAD metrics, assign COIs
6. Apply the logistic regression models to each scenario, using the genes assigned to COIs as control genes

Each simulation scenario was iterated across 1000 random datasets. The performance of the approach was assessed by summarizing the mean LOF, MIS, and SYN and the standard error of those estimates in each simulation scenario.

## Results

### WES variant call and filtering

WES data for all 190 IASPSAD subjects passed quality control measurements, with mean sequencing depth across all samples at 60.6x (standard deviation 12.02), with 96.7% of the target covered at ≥10x. A total of 782,711 variants were detected with 677,758 SNPs and an additional 109,526 insertion/deletions (indels). For quality control, SNPs and indels were excluded if they fell into GATK VQSR tranches 99.0 or greater, indicating that 99% of the true variants present in the sample will be retained in the filtered set (38,503 variants removed), to avoid the rising rates of false positive variant calls at this and more inclusive thresholds. Variants with MAF ≥ 0.05 in gnomAD European (non-Finnish), non-cancer samples were excluded which resulted in a final set of 652,428 variants (91,780 removed) with 2,328 LOF, 31,015 MIS, and 46,046 SYN variants left for exome analysis.

### Description of GOIs

Figure 3 shows the overview of the analysis framework. The correlation between the LOF, MIS, and SYN counts for all genes from gnomAD and the observed counts from the IASPSAD subjects were 0.26, 0.80, and 0.76, respectively. This indicates that at least for MIS and SYN variants, there is sufficiently strong evidence that the gnomAD database information can be utilized to group genes for a case-only within-sample analysis. Table 2 shows the correlation between the 10 gene metrics used for identifying control genes from the gnomAD database, indicating that these annotations are measuring disparate genomic features, which supports the inclusion of all ten metrics in the multivariate clustering. Table 3 describes the resulting trees cut at various heights in the hierarchical clustering analysis. We chose to cut all three GOI set trees at height=0.09 (representing 9% of the tree), a value which achieves a median cluster size of 8 genes, while still only utilizing just under 20% of all genes in the largest COIs. Figure 4 provides a visual representation of one of the branches, representing one cluster of the pruned tree with the GOI (*ADH4*) labeled in red. From a methodological standpoint, the size of GOI sets will depend on the hypothesis and application. Therefore, the decision on cut height will need to be made on an experiment-wise basis in order to balance the need for large clusters (for maximum statistical power for comparisons) and smaller overall proportion of all genes included in the COIs (for tight clusters with high similarity across metrics). Figure 5 illustrates the relationship between GOI set size, median cluster size, and proportion of the exome included in the COI.

**Figure 3:**
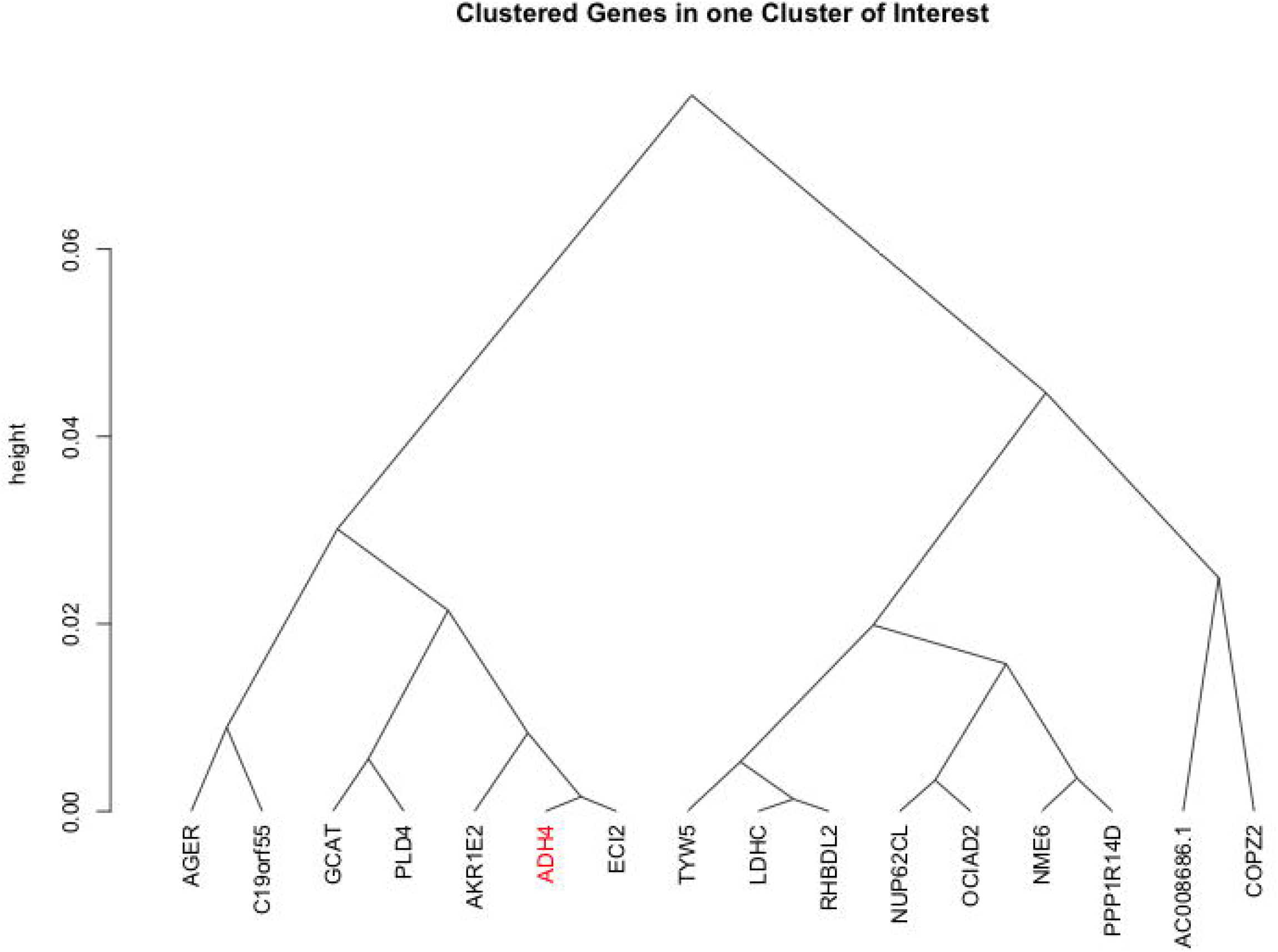
Flow chart demonstrating the analysis framework. (*Figure created with BioRender.com*)

**Figure 4:**
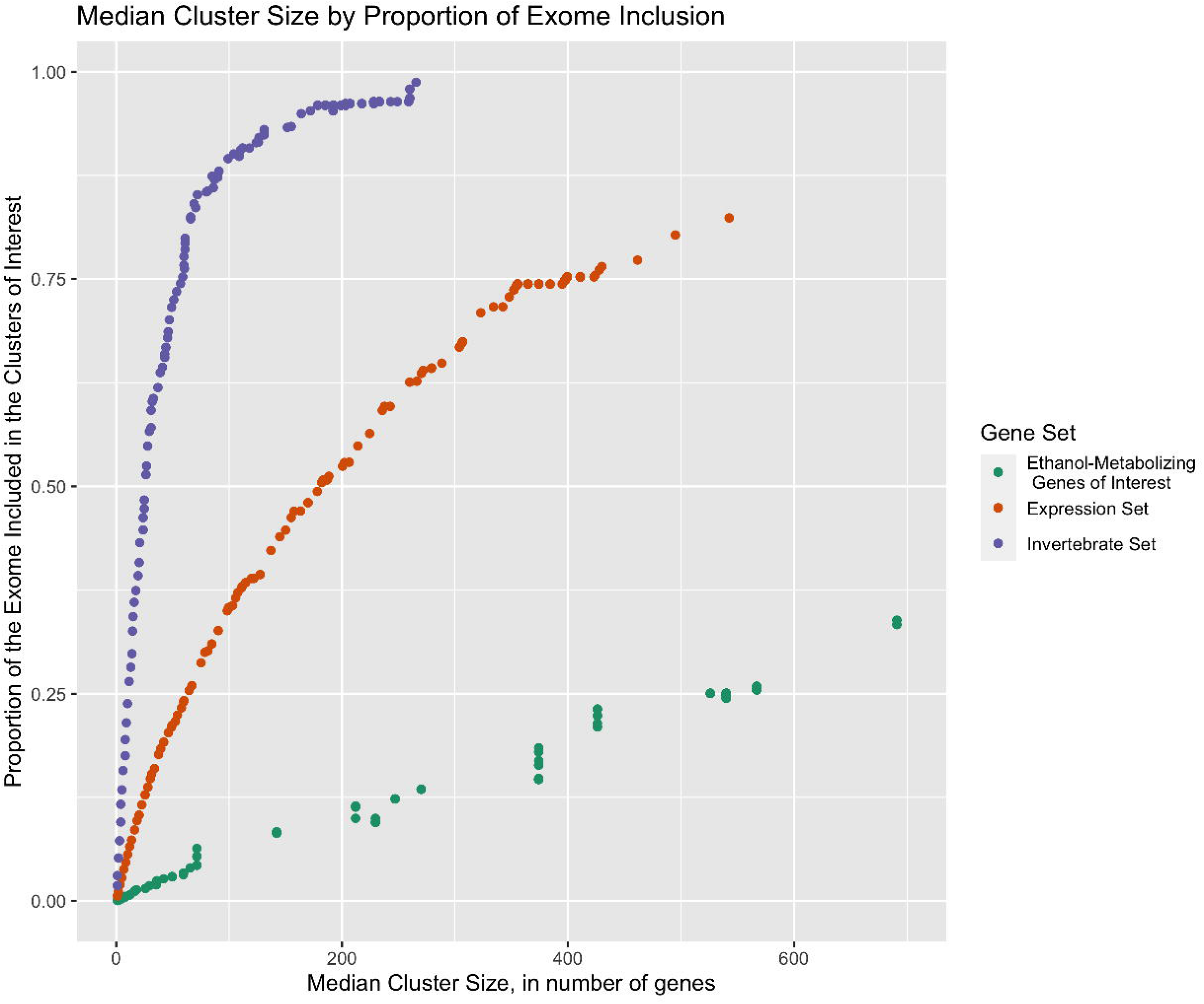
Example of one branch (cluster of interest) from the final, pruned hierarchical clustering tree.

**Figure 5:**
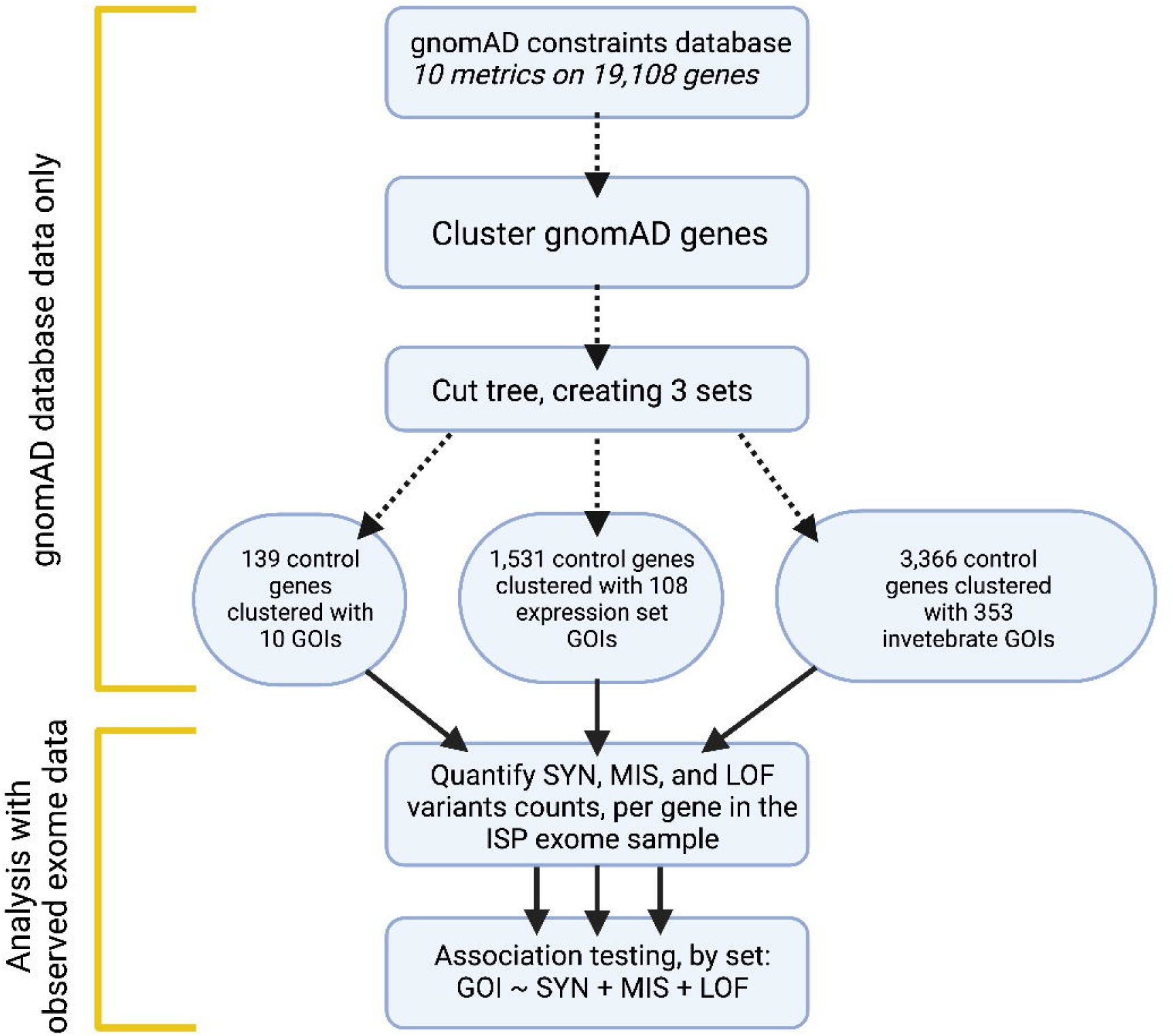
Relationship between median number of genes per cluster of interest and proportion of the exome included in the cluster of interest.

**Table 3:** For each of the three GOI test sets, median cluster size (Med. Size), number of total genes, combined across all clusters of interest (No.Genes), and proportion of the exome retained in the clusters of interest (Prop. Gen) for hierarchical clustering trees cut at various heights.

|  | **Primary human ethanol**<br>**metabolizing GOI set** |  |  | **Including mouse brain**<br>**expression set GOI set** |  |  | **Including invertebrate**<br>**GOI set** |  |  |
| --- | --- | --- | --- | --- | --- | --- | --- | --- | --- |
| **Height** | **Med.**<br>**Size** | **No.**<br>**Genes** | **Prop.** | **Med.**<br>**Size** | **No.**<br>**Genes** | **Prop.** | **Med.**<br>**Size** | **No.**<br>**Genes** | **Prop.** |
| 0 | 1 | 10 | 0.001 | 1 | 117 | 0.006 | 1 | 359 | 0.019 |
| 0.01 | 2.5 | 22 | 0.001 | 2 | 214 | 0.011 | 1 | 589 | 0.031 |
| 0.02 | 3.5 | 35 | 0.002 | 3 | 376 | 0.02 | 2 | 993 | 0.052 |
| 0.03 | 4 | 42 | 0.002 | 4 | 538 | 0.028 | 3 | 1388 | 0.073 |
| 0.04 | 6.5 | 75 | 0.004 | 5 | 731 | 0.038 | 4 | 1822 | 0.095 |
| 0.05 | 7.5 | 83 | 0.004 | 5 | 891 | 0.047 | 4 | 2230 | 0.117 |
| 0.06 | 8 | 97 | 0.005 | 8 | 1072 | 0.056 | 5 | 2561 | 0.134 |
| 0.07 | 8 | 108 | 0.006 | 8 | 1252 | 0.066 | 6 | 3007 | 0.157 |
| 0.08 | 11.5 | 130 | 0.007 | 10 | 1403 | 0.073 | 8 | 3351 | 0.175 |
| .09 | 12.5 | 149 | 0.008 | 11 | 1639 | 0.086 | 8 | 3719 | 0.195 |
| 0.15 | 26 | 290 | 0.015 | 22 | 2818 | 0.147 | 14.5 | 6220 | 0.326 |
| 0.2 | 36 | 470 | 0.025 | 26 | 3662 | 0.192 | 20.5 | 7798 | 0.408 |
| 0.25 | 59.5 | 608 | 0.032 | 34 | 4285 | 0.224 | 25 | 9238 | 0.483 |
| 0.5 | 247 | 2353 | 0.123 | 94 | 8549 | 0.447 | 60 | 14849 | 0.777 |
| 0.75 | 426 | 4272 | 0.224 | 152 | 12227 | 0.64 | 118 | 17349 | 0.908 |
| 1.00 | 19,108 | 19,108 | 1.00 | 19,108 | 19,108 | 1.00 | 19,108 | 19,108 | 1.00 |

Med. Size: Median cluster size
No. Genes: Number of genes across all clusters of interest
Prop. Gen.: Proportion of the exome included

### Simulation results

Full simulation results appear in Supplemental Table S4. In summary, these simulations demonstrated the validity of the logistic regression framework to answer the directed hypotheses regarding counts of LOF, MIS, and SYN variants in GOIs, as compared to matched control genes. The approach was able to accurately identify significant LOF, MIS, and SYN effects where they were simulated to exist in the data. Point estimates were somewhat overestimated in most cases, but for MIS and SYN variants, the true effect fell within 2 standard deviations of the estimated values. Figure 6 shows the estimates, plus 2 standard deviations, as well as the true simulated effect for the LOF (panel a), MIS (panel b), and SYN (panel c) variants across 9 simulation scenarios.

**Figure 6:**
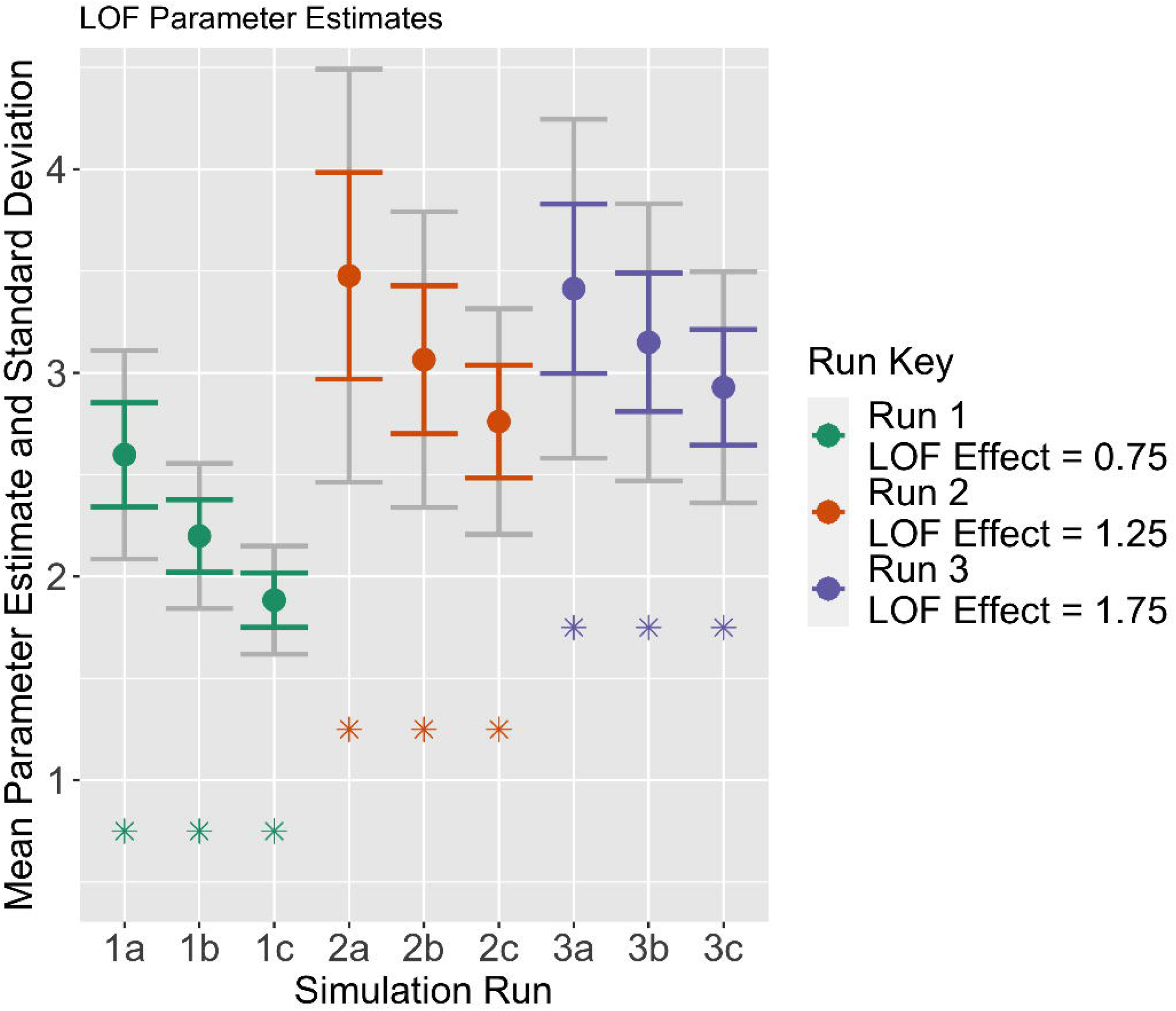
Simulation estimates of parameter values with standard deviations and true simulated values. One standard deviation from the point estimate is shown in the same color as the plotted point with one additional standard deviation shown in gray. The starred point shows the true, simulated point value. Estimates shown for (a) LOF, (b) MIS, and (c) SYN variants.

### Association analysis

For each of the three GOI sets tested, logistic regression was performed where all GOIs served as “cases” and all other genes in the associated COIs served as “controls”. The total set size for the three GOI sets were 149 for the primary human alcohol metabolizing GOI, 1,639 for the mouse PFC brain expression hub GOI, and 3,719 for altered ethanol response invertebrate GOI, respectively. Supplemental Tables S5-S7 list the control genes for each model. Estimates for the logistic regression models are shown in Table 4. Using an alpha cutoff of 0.05, the human ethanol metabolizing GOI did not show a statistically significant difference between the number of LOF, MIS, or SYN variants in GOI compared to control genes. For the mouse PFC brain expression GOI, a significant difference in counts of SYN variants with odds ratio (OR) of 1.21 (*p*-value = 0.0006) was observed, indicating that the addition of a single synonymous variant conferred odds of a gene being a GOI 1.21 times higher than the odds of being a control gene.

**Table 4:** Logistic regression results.

| **Primary Human Ethanol Metabolizing GOI set**<br>N=149 (10 GOI, 139 control genes) |  |  |  |
| --- | --- | --- | --- |
| **Parameter** | **Estimate** | **Standard Error** | **Marginal P-value** |
| *Intercept* | -2.36 | 0.531 | <0.0001 |
| *LOF Alc* | -0.17 | 0.900 | 0.849 |
| *MIS Alc* | -0.064 | 0.206 | 0.757 |
| *SYN Alc* | -0.11 | 0.276 | 0.678 |

| **Mouse Brain Expression GOI Set**<br>N=1,639 (117 GOI, 1,522 control genes) |  |  |  |
| --- | --- | --- | --- |
| *Parameter* | *Estimate* | *Standard Error* | *Marginal P-value* |
| *Intercept* | -2.74 | 0.126 | <0.001 |
| *LOF Alc* | -0.80 | 0.552 | 0.1462 |
| *MIS Alc* | -0.05 | 0.044 | 0.2296 |
| *SYN Alc* | 0.18 | 0.053 | 0.0006 |

| **Invertebrate GOI Set**<br>N=3,719 (359 GOI, 3,361 control genes) |  |  |  |
| --- | --- | --- | --- |
| *Parameter* | *Estimate* | *Standard Error* | *Marginal P-value* |
| *Intercept* | -2.34 | 0.071 | <0.001 |
| *LOF Alc* | *-0.10* | *0.166* | *0.5330* |
| *MIS Alc* | *0.004* | *0.022* | *0.8367* |
| *SYN Alc* | *0.06* | *0.026* | *0.0169* |

Finally, for the altered ethanol response invertebrate GOI, a significant difference in the number of SYN variants with an OR of 1.06 was observed (*p*-value 0.0169).

## Discussion

In this study, we sought to perform a hypothesis driven case-only exome analysis in severe AD cases using three GOI sets with prior evidence for ethanol metabolism in humans, or implicated in alcohol phenotypes in model organism studies in mouse and invertebrates. While we observed differences in the number of SYN variants for mouse and invertebrate GOIs, the results do not support the hypothesis that ethanol metabolizing genes, in particular those directly involved in humans, are largely depleted for LOF variants in severe AD cases. While MIS or LOF variants have more direct effect on gene products, SYN mutations can impact the speed of messenger RNA (mRNA) translation processes by changing the codon to one with different transfer RNA (tRNA) abundance or by altering folding and stability of mRNA, producing secondary structures that are less efficiently recognized for mRNA processing(50). In particular, recent oncological findings suggest that SYN mutations might play a role in the development of cancer by altering codon optimization and translational velocity. However, SYN mutations are still largely considered to be functionally silent.

Genetic studies of cardiac disease(51–53), obesity(54), Alzheimer disease(55), and nonalcoholic fatty liver disease(56) consistently show that MIS and LOF alleles are common in the human genome, alter disease risk, and in some cases are protective(57,58). Such protective effects of functional variants are well documented for alcohol-related phenotypes. Most recently, rs75967634 in *ADH1B* was also found to be associated with problematic alcohol use in individuals of European ancestry(21). Although of low frequency outside of east Asia,

*ADH1B*2* (rs1229984) is associated with both AUD diagnosis(34,59) and the problem drinking component of the AUDIT(20) in European populations. Together, these results suggest that although effects of variants in ethanol metabolizing enzymes are well documented across different populations, our findings based on our modestly sized sample of severe AD cases does not provide evidence in support of a significant depletion of LOF or MIS variation in these genes.

In this study, we presented a framework for analyzing case-only exome variation data in the absence of appropriate control subjects. It is not uncommon in genetic studies to obtain molecular data on a set of subjects who are all positive for a dichotomous phenotype. Therefore, methods have been developed for intentional case-only study designs, in particular for gene-by-environment interaction studies(60) or polygenic risk scores(61). However, current case-only analysis frameworks generally consider an environmental exposure as the dichotomous outcome being tested for association with genetic variation within a sample of all cases, and offer no information regarding phenotypic variation due to the primary disease state. Furthermore, when controls have not been genotyped or sequenced alongside cases in the same study, publicly available datasets may also provide appropriate population controls with careful ancestral matching. Additionally, methods such as the Robust Variance Score Statistic (RVS) method(62), or the burden test implemented in the TASER software(63) which is an extension of RVS method that offers improved adjustment for sample differences in case/control analyses that can be used in such cases. Other methods such as iECAT(64) does not require individual-level genotype data from population controls, but rather conducts an adjusted association testing using only allele counts. However, due to differences in sampling techniques or technology, filtering criteria, and variant calling, using these methods may not always be feasible. Importantly, none of these methods offers correction or adjustment for related samples such as the IASPSAD sample analyzed in this study. We therefore sought to model case/control status of individual genes within a case only sample of severely affected subjects with AD cases using a novel design. In contrast to studies that use external control subjects, our proposed framework used individual cases’ exome data to assign case status to GOI and control status to matched genes as described in the methods section. This strategy removes the need for careful correction or adjustment for subtle population structure in exome studies, and further leverages external information from large sequencing studies such as gnomAD to ensure gene matching is robust by using a multivariate agnostic hierarchical clustering approach. Therefore, this methodological framework represents an interpretable, straight-forward, and computationally affordable approach that is easily implemented using existing software and tools. We additionally note that this framework can also be extended to model quantitative measures within cases, such as maximum number of drinks per day with minimal adjustment.

Finally, our simulation studies demonstrated that while the approach has sufficient power to detect real effects, the estimates of these effects may show some inflation under certain distributional conditions. It is recommended that future applications of this approach utilize some simulations to determine empirically expectations of power and identify potential sources of bias *a priori*. Conservatively, the simulation results indicate that the significant estimate of the effect of synonymous variants in the mouse expression set is unlikely attributable to positive bias alone. Furthermore, while such exploration was beyond the scope of the current study, different experimental designs may warrant further simulations to empirically guide the choice of tree cutpoints to balance cluster size and proportion of the exome included in the COIs.

The findings presented in this study should be interpreted in the context of three important limitations. First, our modest sample size of 190 affected subjects with exome data is limited, and the tests conducted on each GOI set may not be sufficiently powered to detect significant differences. Second, while we empirically modeled the median number of matched genes in each cluster against the proportion of the genome in the final clustered set to choose an appropriate pruning parameter, simulations to evaluate the impact of various parameter choices such as tree pruning height, or proportion of the genome included in gene clusters could provide better benchmarking for selecting these parameters in future studies. Third, as more exome data from ancestrally diverse populations become available, future analyses could attempt to replicate these findings in larger, more diverse samples to improve the generalizability of these findings and the utility of our case-only exome analysis framework in other populations and phenotypes.

## Acknowledgements

This work was supported in part by the Virginia Commonwealth University Alcohol Research Center and NIAAA P50AA022537.

## Supplemental Information Captions

**Supplemental Table S1:** The genes in the expression set (n=109 genes), with annotation indicating which were present in gnomAD (108), which were present in the invertebrate set (108), and which were present in the human GOI set (1).

**Supplemental Table S2:** The list of invertebrate genes (n = 358 genes) with annotation indicating which were present in gnomAD (353), which were present in the expression set (108), and which were present in the human GOI set (11).

**Supplemental Section S1: Simulation Details**

**Supplemental Table S3:** Parameters used to generate random data for the simulations.

**Supplemental Table S4:** Parameter estimates from simulations.

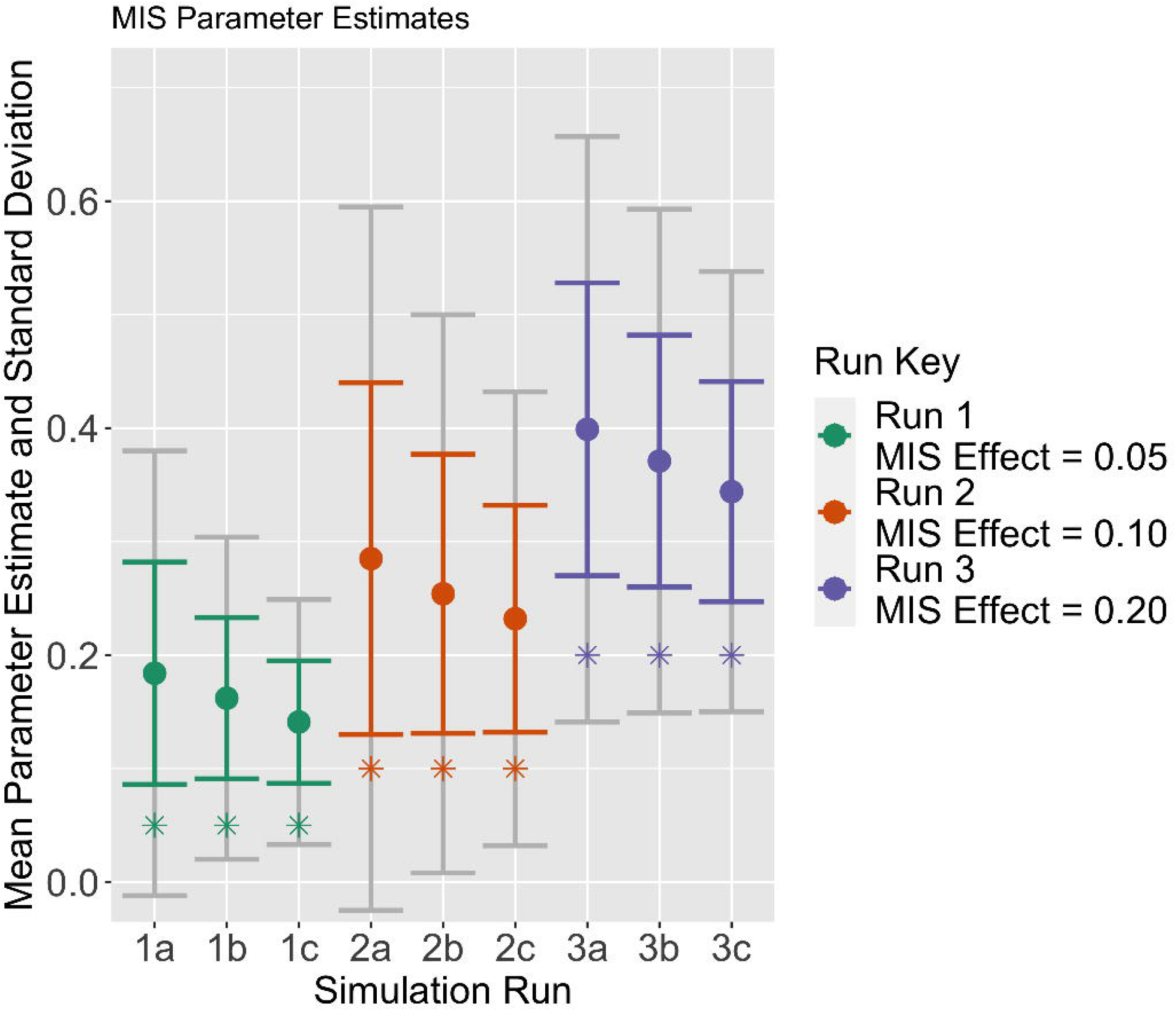

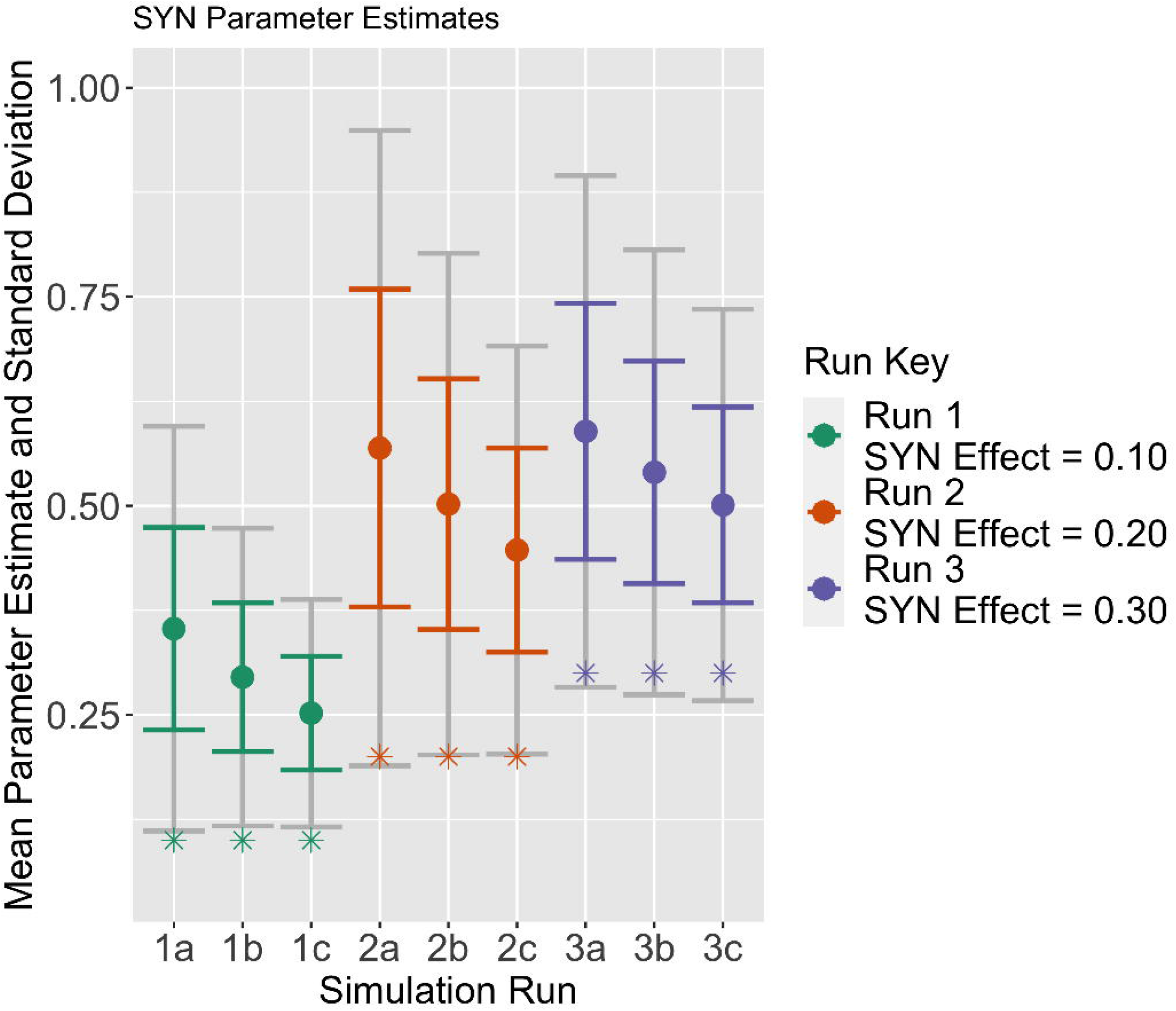

