## Supplemental Material for "Case-only exome variation analysis of severe alcohol dependence using a multivariate hierarchical gene clustering approach"

**TITLE**

*Currently at Veterinary Information Network

Supplement Contents:

Supplemental Table 1
Supplemental Table 2
Supplemental Section 1
Supplemental Table 3
Supplemental Table 4

**Supplemental Table S1:** The genes in the expression set (n=109 genes), with annotation indicating which were present in gnomAD (108), which were present in the invertebrate set (108), and which were present in the human GOI set (1).

| Gene | Transcript | Present in gnomAD | Present in invertebrate set | Present in primary GOI set |
|---|---|---|---|---|
| ACTL6A | ENST00000429709 | TRUE | TRUE | FALSE |
| ACTL6B | ENST00000160382 | TRUE | TRUE | FALSE |
| ADCY1 | ENST00000297323 | TRUE | TRUE | FALSE |
| AKT1 | ENST00000554581 | TRUE | TRUE | FALSE |
| AKT3 | ENST00000366539 | TRUE | TRUE | FALSE |
| ALDH2 | ENST00000261733 | TRUE | TRUE | TRUE |
| ALDH6A1 | ENST00000553458 | TRUE | TRUE | FALSE |
| ALDH9A1 | ENST00000354775 | TRUE | TRUE | FALSE |
| ARF6 | ENST00000298316 | TRUE | TRUE | FALSE |
| ARL6IP5 | ENST00000273258 | TRUE | TRUE | FALSE |
| ATP1A1 | ENST00000537345 | TRUE | TRUE | FALSE |
| ATP1A3 | ENST00000545399 | TRUE | TRUE | FALSE |
| AUTS2 | ENST00000342771 | TRUE | TRUE | FALSE |
| CDC42 | ENST00000344548 | TRUE | TRUE | FALSE |
| CHP1 | ENST00000334660 | TRUE | TRUE | FALSE |
| CHRNA6 | ENST00000276410 | TRUE | TRUE | FALSE |
| CLIC4 | ENST00000374379 | TRUE | TRUE | FALSE |
| COX6C | ENST00000520468 | TRUE | TRUE | FALSE |
| CRYAB | ENST00000533475 | TRUE | TRUE | FALSE |
| CSAD | ENST00000267085 | TRUE | TRUE | FALSE |
| CSNK1A1 | ENST00000515768 | TRUE | TRUE | FALSE |
| CSNK1D | ENST00000314028 | TRUE | TRUE | FALSE |
| CSNK1E | ENST00000396832 | TRUE | TRUE | FALSE |
| DLG1 | ENST00000346964 | TRUE | TRUE | FALSE |
| DLG2 | ENST00000376104 | TRUE | TRUE | FALSE |
| DLG4 | ENST00000399510 | TRUE | TRUE | FALSE |
| DNM1 | ENST00000372923 | TRUE | TRUE | FALSE |
| DNM2 | ENST00000389253 | TRUE | TRUE | FALSE |
| DNM3 | ENST00000358155 | TRUE | TRUE | FALSE |
| DUSP10 | ENST00000366899 | TRUE | TRUE | FALSE |
| EPS8 | ENST00000281172 | TRUE | TRUE | FALSE |
| FADS1 | ENST00000350997 | TRUE | TRUE | FALSE |
| FADS2 | ENST00000278840 | TRUE | TRUE | FALSE |
| FGFR1 | ENST00000425967 | TRUE | TRUE | FALSE |
| FGFR2 | ENST00000457416 | TRUE | TRUE | FALSE |

| | | | | |
|---|---|---|---|---|
| FGFR3 | ENST00000340107 | TRUE | TRUE | FALSE |
| FOXO3 | ENST00000406360 | TRUE | TRUE | FALSE |
| FSTL3 | ENST00000166139 | TRUE | TRUE | FALSE |
| GABBR1 | ENST00000377034 | TRUE | TRUE | FALSE |
| GAD1 | ENST00000358196 | TRUE | TRUE | FALSE |
| GAD2 | ENST00000376261 | TRUE | TRUE | FALSE |
| GPC6 | ENST00000377047 | TRUE | TRUE | FALSE |
| GRIN1 | ENST00000371553 | TRUE | TRUE | FALSE |
| HOMER1 | ENST00000334082 | TRUE | TRUE | FALSE |
| HOMER2 | ENST00000304231 | TRUE | TRUE | FALSE |
| HOMER3 | ENST00000539827 | TRUE | TRUE | FALSE |
| IGF1R | ENST00000268035 | TRUE | TRUE | FALSE |
| IRS4 | ENST00000372129 | TRUE | TRUE | FALSE |
| ITGB1 | ENST00000396033 | TRUE | TRUE | FALSE |
| ITGB2 | ENST00000397850 | TRUE | TRUE | FALSE |
| ITGB5 | ENST00000296181 | TRUE | TRUE | FALSE |
| KCNMA1 | ENST00000404857 | TRUE | TRUE | FALSE |
| KCNQ2 | ENST00000359125 | TRUE | TRUE | FALSE |
| KCNQ5 | ENST00000342056 | TRUE | TRUE | FALSE |
| MAP4K5 | ENST00000013125 | TRUE | TRUE | FALSE |
| MAPK1 | ENST00000215832 | TRUE | TRUE | FALSE |
| MAPK10 | ENST00000359221 | TRUE | TRUE | FALSE |
| MAPK8 | ENST00000374189 | TRUE | TRUE | FALSE |
| MAPK9 | ENST00000452135 | TRUE | TRUE | FALSE |
| MARK1 | ENST00000366917 | TRUE | TRUE | FALSE |
| MARK2 | ENST00000402010 | TRUE | TRUE | FALSE |
| MARK4 | ENST00000262891 | TRUE | TRUE | FALSE |
| NCAM1 | ENST00000524665 | TRUE | TRUE | FALSE |
| NCAM2 | ENST00000400546 | TRUE | TRUE | FALSE |
| NPY | ENST00000407573 | TRUE | TRUE | FALSE |
| PBRM1 | ENST00000394830 | TRUE | TRUE | FALSE |
| PCSK2 | ENST00000262545 | TRUE | TRUE | FALSE |
| PDPK1 | ENST00000342085 | TRUE | TRUE | FALSE |
| PER1 | ENST00000317276 | TRUE | TRUE | FALSE |
| PER2 | ENST00000254657 | TRUE | TRUE | FALSE |
| PIK3CA | ENST00000263967 | TRUE | TRUE | FALSE |
| PIK3R1 | ENST00000521381 | TRUE | TRUE | FALSE |
| PPIA | ENST00000468812 | TRUE | TRUE | FALSE |
| PRKACA | ENST00000308677 | TRUE | TRUE | FALSE |
| PRKACB | ENST00000370685 | TRUE | TRUE | FALSE |

| | | | | |
|---|---|---|---|---|
| PRKAR2A | ENST00000265563 | TRUE | TRUE | FALSE |
| PRKAR2B | ENST00000265717 | TRUE | TRUE | FALSE |
| PTEN | ENST00000371953 | TRUE | TRUE | FALSE |
| RAB3C | ENST00000282878 | TRUE | TRUE | FALSE |
| RAC1 | ENST00000356142 | TRUE | TRUE | FALSE |
| RAC2 | ENST00000249071 | TRUE | TRUE | FALSE |
| RFNG | ENST00000310496 | TRUE | TRUE | FALSE |
| RHEBL1 | ENST00000301068 | TRUE | TRUE | FALSE |
| RHOA | ENST00000418115 | TRUE | TRUE | FALSE |
| SDC1 | ENST00000381150 | TRUE | TRUE | FALSE |
| SDC2 | ENST00000302190 | TRUE | TRUE | FALSE |
| SDC4 | ENST00000372733 | TRUE | TRUE | FALSE |
| SGK1 | ENST00000367858 | TRUE | TRUE | FALSE |
| SLC18A2 | ENST00000298472 | TRUE | TRUE | FALSE |
| SLC32A1 | ENST00000217420 | TRUE | TRUE | FALSE |
| SLC9A3R1 | ENST00000262613 | TRUE | TRUE | FALSE |
| SLC9A3R2 | ENST00000424542 | TRUE | TRUE | FALSE |
| SMARCA2 | ENST00000382203 | TRUE | TRUE | FALSE |
| SMARCA4 | ENST00000429416 | TRUE | TRUE | FALSE |
| SMARCC1 | ENST00000254480 | TRUE | TRUE | FALSE |
| SMARCD1 | ENST00000394963 | TRUE | TRUE | FALSE |
| SMARCD2 | ENST00000448276 | TRUE | TRUE | FALSE |
| SMARCD3 | ENST00000262188 | TRUE | TRUE | FALSE |
| SREBF1 | ENST00000355815 | TRUE | TRUE | FALSE |
| SREBF2 | ENST00000361204 | TRUE | TRUE | FALSE |
| STX1A | ENST00000222812 | TRUE | TRUE | FALSE |
| STX1B | ENST00000215095 | TRUE | TRUE | FALSE |
| STX2 | ENST00000392373 | TRUE | TRUE | FALSE |
| STX3 | ENST00000337979 | TRUE | TRUE | FALSE |
| STXBP1 | ENST00000373302 | TRUE | TRUE | FALSE |
| SYN2 | NA | FALSE | TRUE | FALSE |
| TAOK1 | ENST00000261716 | TRUE | TRUE | FALSE |
| TH | ENST00000381178 | TRUE | TRUE | FALSE |
| TPH2 | ENST00000333850 | TRUE | TRUE | FALSE |

**Supplemental Table S2:** The list of invertebrate genes (n = 358 genes) with annotation indicating which were present in gnomAD (353), which were present in the expression set (108), and which were present in the human GOI set (11).

| Gene | Transcript | Present in gnomAD | Present in expression set | Present in primary GOI set |
|---|---|---|---|---|
| ABCG2 | ENST00000237612 | TRUE | FALSE | FALSE |
| ACAA2 | ENST00000285093 | TRUE | FALSE | FALSE |
| ACADM | ENST00000420607 | TRUE | FALSE | FALSE |
| ACADSB | ENST00000358776 | TRUE | FALSE | FALSE |
| ACSL5 | ENST00000356116 | TRUE | FALSE | FALSE |
| ACSS1 | ENST00000323482 | TRUE | FALSE | FALSE |
| ACSS2 | ENST00000253382 | TRUE | FALSE | FALSE |
| ACTL6A | ENST00000429709 | TRUE | TRUE | FALSE |
| ACTL6B | ENST00000160382 | TRUE | TRUE | FALSE |
| ADCY1 | ENST00000297323 | TRUE | TRUE | FALSE |
| ADH1A | ENST00000209668 | TRUE | FALSE | TRUE |
| ADH1B | ENST00000305046 | TRUE | FALSE | TRUE |
| ADH1C | NA | FALSE | FALSE | TRUE |
| AKT1 | ENST00000554581 | TRUE | TRUE | FALSE |
| AKT2 | ENST00000392038 | TRUE | FALSE | FALSE |
| AKT3 | ENST00000366539 | TRUE | TRUE | FALSE |
| ALDH16A1 | ENST00000293350 | TRUE | FALSE | FALSE |
| ALDH18A1 | ENST00000371224 | TRUE | FALSE | FALSE |
| ALDH1A1 | ENST00000297785 | TRUE | FALSE | TRUE |
| ALDH1A2 | ENST00000249750 | TRUE | FALSE | FALSE |
| ALDH1A3 | ENST00000329841 | TRUE | FALSE | FALSE |
| ALDH1B1 | ENST00000377698 | TRUE | FALSE | FALSE |
| ALDH1L1 | ENST00000273450 | TRUE | FALSE | FALSE |
| ALDH1L2 | ENST00000258494 | TRUE | FALSE | FALSE |
| ALDH2 | ENST00000261733 | TRUE | TRUE | TRUE |
| ALDH3A1 | ENST00000457500 | TRUE | FALSE | FALSE |
| ALDH3A2 | ENST00000339618 | TRUE | FALSE | FALSE |
| ALDH3B1 | ENST00000539229 | TRUE | FALSE | FALSE |
| ALDH3B2 | ENST00000349015 | TRUE | FALSE | FALSE |
| ALDH4A1 | ENST00000375341 | TRUE | FALSE | FALSE |
| ALDH5A1 | ENST00000348925 | TRUE | FALSE | FALSE |
| ALDH6A1 | ENST00000553458 | TRUE | TRUE | FALSE |
| ALDH7A1 | ENST00000409134 | TRUE | FALSE | FALSE |
| ALDH8A1 | ENST00000265605 | TRUE | FALSE | FALSE |
| ALDH9A1 | ENST00000354775 | TRUE | TRUE | FALSE |
| ALK | ENST00000389048 | TRUE | FALSE | FALSE |
| ARF6 | ENST00000298316 | TRUE | TRUE | FALSE |
| ARFIP1 | ENST00000451320 | TRUE | FALSE | FALSE |
| ARFIP2 | ENST00000254584 | TRUE | FALSE | FALSE |
| ARID1A | ENST00000324856 | TRUE | FALSE | FALSE |
| ARID1B | ENST00000346085 | TRUE | FALSE | FALSE |
| ARID2 | ENST00000334344 | TRUE | FALSE | FALSE |
| ARL6IP5 | ENST00000273258 | TRUE | TRUE | FALSE |
| ARNTL | ENST00000389707 | TRUE | FALSE | FALSE |
| ARNTL2 | ENST00000266503 | TRUE | FALSE | FALSE |
| ATP12A | ENST00000218548 | TRUE | FALSE | FALSE |
| ATP1A1 | ENST00000537345 | TRUE | TRUE | FALSE |
| ATP1A2 | ENST00000361216 | TRUE | FALSE | FALSE |
| ATP1A3 | ENST00000545399 | TRUE | TRUE | FALSE |
| ATP1A4 | ENST00000368081 | TRUE | FALSE | FALSE |
| ATP4A | ENST00000262623 | TRUE | FALSE | FALSE |
| AUTS2 | ENST00000342771 | TRUE | TRUE | FALSE |
| BBS1 | ENST00000318312 | TRUE | FALSE | FALSE |
| BRD7 | ENST00000394689 | TRUE | FALSE | FALSE |
| BRD9 | ENST00000467963 | TRUE | FALSE | FALSE |

| | | | | |
|---|---|---|---|---|
| CASK | ENST00000378166 | TRUE | FALSE | FALSE |
| CBS | ENST00000398165 | TRUE | FALSE | FALSE |
| CBSL | NA | FALSE | FALSE | FALSE |
| CD180 | ENST00000256447 | TRUE | FALSE | FALSE |
| CDC42 | ENST00000344548 | TRUE | TRUE | FALSE |
| CHAT | ENST00000337653 | TRUE | FALSE | FALSE |
| CHP1 | ENST00000334660 | TRUE | TRUE | FALSE |
| CHP2 | ENST00000300113 | TRUE | FALSE | FALSE |
| CHRNA1 | ENST00000261007 | TRUE | FALSE | FALSE |
| CHRNA2 | ENST00000407991 | TRUE | FALSE | FALSE |
| CHRNA3 | ENST00000326828 | TRUE | FALSE | FALSE |
| CHRNA4 | ENST00000370263 | TRUE | FALSE | FALSE |
| CHRNA6 | ENST00000276410 | TRUE | TRUE | FALSE |
| CHRNB2 | ENST00000368476 | TRUE | FALSE | FALSE |
| CHRNB4 | ENST00000261751 | TRUE | FALSE | FALSE |
| CLIC1 | ENST00000375780 | TRUE | FALSE | FALSE |
| CLIC2 | ENST00000369449 | TRUE | FALSE | FALSE |
| CLIC3 | ENST00000494426 | TRUE | FALSE | FALSE |
| CLIC4 | ENST00000374379 | TRUE | TRUE | FALSE |
| CLIC5 | ENST00000185206 | TRUE | FALSE | FALSE |
| CLIC6 | ENST00000349499 | TRUE | FALSE | FALSE |
| COX6C | ENST00000520468 | TRUE | TRUE | FALSE |
| CPT1B | ENST00000360719 | TRUE | FALSE | FALSE |
| CRYAA | ENST00000291554 | TRUE | FALSE | FALSE |
| CRYAB | ENST00000533475 | TRUE | TRUE | FALSE |
| CSAD | ENST00000267085 | TRUE | TRUE | FALSE |
| CSNK1A1 | ENST00000515768 | TRUE | TRUE | FALSE |
| CSNK1D | ENST00000314028 | TRUE | TRUE | FALSE |
| CSNK1E | ENST00000396832 | TRUE | TRUE | FALSE |
| CTBP1 | ENST00000290921 | TRUE | FALSE | FALSE |
| CTBP2 | ENST00000309035 | TRUE | FALSE | FALSE |
| CTSH | ENST00000220166 | TRUE | FALSE | FALSE |
| CTSK | ENST00000271651 | TRUE | FALSE | FALSE |
| CTSL | ENST00000343150 | TRUE | FALSE | FALSE |
| CTSS | ENST00000368985 | TRUE | FALSE | FALSE |
| CTSV | ENST00000259470 | TRUE | FALSE | FALSE |
| DBH | ENST00000393056 | TRUE | FALSE | FALSE |
| DDC | ENST00000444124 | TRUE | FALSE | FALSE |
| DGKQ | ENST00000273814 | TRUE | FALSE | FALSE |
| DLG1 | ENST00000346964 | TRUE | TRUE | FALSE |
| DLG2 | ENST00000376104 | TRUE | TRUE | FALSE |
| DLG3 | ENST00000374360 | TRUE | FALSE | FALSE |
| DLG4 | ENST00000399510 | TRUE | TRUE | FALSE |
| DNM1 | ENST00000372923 | TRUE | TRUE | FALSE |
| DNM2 | ENST00000389253 | TRUE | TRUE | FALSE |
| DNM3 | ENST00000358155 | TRUE | TRUE | FALSE |
| DPF1 | ENST00000355526 | TRUE | FALSE | FALSE |
| DPF2 | ENST00000528416 | TRUE | FALSE | FALSE |
| DPF3 | ENST00000541685 | TRUE | FALSE | FALSE |
| DRD1 | ENST00000393752 | TRUE | FALSE | FALSE |
| DRD5 | ENST00000304374 | TRUE | FALSE | FALSE |
| DUSP10 | ENST00000366899 | TRUE | TRUE | FALSE |
| EGFR | ENST00000275493 | TRUE | FALSE | FALSE |
| EI24 | ENST00000278903 | TRUE | FALSE | FALSE |
| EPS8 | ENST00000281172 | TRUE | TRUE | FALSE |
| EPS8L1 | ENST00000201647 | TRUE | FALSE | FALSE |
| EPS8L2 | ENST00000533256 | TRUE | FALSE | FALSE |
| EPS8L3 | ENST00000369805 | TRUE | FALSE | FALSE |
| ERBB2 | ENST00000269571 | TRUE | FALSE | FALSE |
| ERBB3 | ENST00000267101 | TRUE | FALSE | FALSE |
| ERBB4 | ENST00000342788 | TRUE | FALSE | FALSE |
| FADS1 | ENST00000350997 | TRUE | TRUE | FALSE |

| | | | | |
|---|---|---|---|---|
| FADS2 | ENST00000278840 | TRUE | TRUE | FALSE |
| FADS3 | ENST00000278829 | TRUE | FALSE | FALSE |
| FBRSL1 | ENST00000434748 | TRUE | FALSE | FALSE |
| FGFR1 | ENST00000425967 | TRUE | TRUE | FALSE |
| FGFR2 | ENST00000457416 | TRUE | TRUE | FALSE |
| FGFR3 | ENST00000340107 | TRUE | TRUE | FALSE |
| FGFR4 | ENST00000292408 | TRUE | FALSE | FALSE |
| FOXO1 | ENST00000379561 | TRUE | FALSE | FALSE |
| FOXO3 | ENST00000406360 | TRUE | TRUE | FALSE |
| FOXO4 | ENST00000374259 | TRUE | FALSE | FALSE |
| FOXO6 | ENST00000372591 | TRUE | FALSE | FALSE |
| FST | ENST00000256759 | TRUE | FALSE | FALSE |
| FSTL3 | ENST00000166139 | TRUE | TRUE | FALSE |
| GABBR1 | ENST00000377034 | TRUE | TRUE | FALSE |
| GAD1 | ENST00000358196 | TRUE | TRUE | FALSE |
| GAD2 | ENST00000376261 | TRUE | TRUE | FALSE |
| GADL1 | ENST00000282538 | TRUE | FALSE | FALSE |
| GFI1 | ENST00000370332 | TRUE | FALSE | FALSE |
| GFI1B | ENST00000339463 | TRUE | FALSE | FALSE |
| GPC1 | ENST00000264039 | TRUE | FALSE | FALSE |
| GPC2 | ENST00000292377 | TRUE | FALSE | FALSE |
| GPC3 | ENST00000394299 | TRUE | FALSE | FALSE |
| GPC4 | ENST00000370828 | TRUE | FALSE | FALSE |
| GPC5 | ENST00000377067 | TRUE | FALSE | FALSE |
| GPC6 | ENST00000377047 | TRUE | TRUE | FALSE |
| GPR21 | ENST00000373642 | TRUE | FALSE | FALSE |
| GPR52 | ENST00000367685 | TRUE | FALSE | FALSE |
| GPR84 | ENST00000551809 | TRUE | FALSE | FALSE |
| GRIN1 | ENST00000371553 | TRUE | TRUE | FALSE |
| GUCY1B3 | ENST00000264424 | TRUE | FALSE | FALSE |
| HADHA | ENST00000380649 | TRUE | FALSE | FALSE |
| HDC | ENST00000267845 | TRUE | FALSE | FALSE |
| HNF4A | ENST00000316099 | TRUE | FALSE | FALSE |
| HNF4G | ENST00000396423 | TRUE | FALSE | FALSE |
| HOMER1 | ENST00000334082 | TRUE | TRUE | FALSE |
| HOMER2 | ENST00000304231 | TRUE | TRUE | FALSE |
| HOMER3 | ENST00000539827 | TRUE | TRUE | FALSE |
| HPGD | ENST00000296522 | TRUE | FALSE | FALSE |
| IGF1R | ENST00000268035 | TRUE | TRUE | FALSE |
| INSR | ENST00000302850 | TRUE | FALSE | FALSE |
| INSRR | ENST00000368195 | TRUE | FALSE | FALSE |
| IRAK1 | ENST00000369980 | TRUE | FALSE | FALSE |
| IRAK2 | ENST00000256458 | TRUE | FALSE | FALSE |
| IRAK3 | ENST00000261233 | TRUE | FALSE | FALSE |
| IRAK4 | ENST00000448290 | TRUE | FALSE | FALSE |
| IRS1 | ENST00000305123 | TRUE | FALSE | FALSE |
| IRS2 | ENST00000375856 | TRUE | FALSE | FALSE |
| IRS4 | ENST00000372129 | TRUE | TRUE | FALSE |
| ITGB1 | ENST00000396033 | TRUE | TRUE | FALSE |
| ITGB2 | ENST00000397850 | TRUE | TRUE | FALSE |
| ITGB3 | ENST00000559488 | TRUE | FALSE | FALSE |
| ITGB5 | ENST00000296181 | TRUE | TRUE | FALSE |
| ITGB7 | ENST00000267082 | TRUE | FALSE | FALSE |
| KCNMA1 | ENST00000404857 | TRUE | TRUE | FALSE |
| KCNQ1 | ENST00000155840 | TRUE | FALSE | FALSE |
| KCNQ2 | ENST00000359125 | TRUE | TRUE | FALSE |
| KCNQ3 | ENST00000388996 | TRUE | FALSE | FALSE |
| KCNQ4 | ENST00000347132 | TRUE | FALSE | FALSE |
| KCNQ5 | ENST00000342056 | TRUE | TRUE | FALSE |
| KCNU1 | ENST00000399881 | TRUE | FALSE | FALSE |
| LFNG | ENST00000222725 | TRUE | FALSE | FALSE |
| LMO1 | ENST00000335790 | TRUE | FALSE | FALSE |

| | | | | |
|---|---|---|---|---|
| LMO2 | ENST00000257818 | TRUE | FALSE | FALSE |
| LMO3 | ENST00000540445 | TRUE | FALSE | FALSE |
| LTK | ENST00000263800 | TRUE | FALSE | FALSE |
| MADD | ENST00000311027 | TRUE | FALSE | FALSE |
| MAP4K1 | ENST00000591517 | TRUE | FALSE | FALSE |
| MAP4K2 | ENST00000294066 | TRUE | FALSE | FALSE |
| MAP4K3 | ENST00000263881 | TRUE | FALSE | FALSE |
| MAP4K5 | ENST00000013125 | TRUE | TRUE | FALSE |
| MAPK1 | ENST00000215832 | TRUE | TRUE | FALSE |
| MAPK10 | ENST00000359221 | TRUE | TRUE | FALSE |
| MAPK3 | ENST00000263025 | TRUE | FALSE | FALSE |
| MAPK8 | ENST00000374189 | TRUE | TRUE | FALSE |
| MAPK9 | ENST00000452135 | TRUE | TRUE | FALSE |
| MARK1 | ENST00000366917 | TRUE | TRUE | FALSE |
| MARK2 | ENST00000402010 | TRUE | TRUE | FALSE |
| MARK3 | ENST00000429436 | TRUE | FALSE | FALSE |
| MARK4 | ENST00000262891 | TRUE | TRUE | FALSE |
| ME1 | ENST00000369705 | TRUE | FALSE | FALSE |
| ME2 | ENST00000321341 | TRUE | FALSE | FALSE |
| ME3 | ENST00000543262 | TRUE | FALSE | FALSE |
| MEF2A | ENST00000354410 | TRUE | FALSE | FALSE |
| MEF2C | ENST00000340208 | TRUE | FALSE | FALSE |
| MEF2D | ENST00000348159 | TRUE | FALSE | FALSE |
| MFNG | ENST00000356998 | TRUE | FALSE | FALSE |
| MOXD1 | ENST00000367963 | TRUE | FALSE | FALSE |
| MTNR1A | ENST00000307161 | TRUE | FALSE | FALSE |
| MTNR1B | ENST00000257068 | TRUE | FALSE | FALSE |
| MYD88 | ENST00000417037 | TRUE | FALSE | FALSE |
| MYL1 | ENST00000352451 | TRUE | FALSE | FALSE |
| MYL3 | ENST00000395869 | TRUE | FALSE | FALSE |
| MYL4 | ENST00000354968 | TRUE | FALSE | FALSE |
| MYL6 | ENST00000550697 | TRUE | FALSE | FALSE |
| MYL6B | ENST00000553066 | TRUE | FALSE | FALSE |
| NALCN | ENST00000251127 | TRUE | FALSE | FALSE |
| NAT10 | ENST00000257829 | TRUE | FALSE | FALSE |
| NCAM1 | ENST00000524665 | TRUE | TRUE | FALSE |
| NCAM2 | ENST00000400546 | TRUE | TRUE | FALSE |
| NFKB1 | ENST00000226574 | TRUE | FALSE | FALSE |
| NFKB2 | ENST00000369966 | TRUE | FALSE | FALSE |
| NFKBIA | ENST00000216797 | TRUE | FALSE | FALSE |
| NFKBIB | ENST00000313582 | TRUE | FALSE | FALSE |
| NOS1 | ENST00000338101 | TRUE | FALSE | FALSE |
| NOS2 | ENST00000313735 | TRUE | FALSE | FALSE |
| NOS3 | ENST00000297494 | TRUE | FALSE | FALSE |
| NPY | ENST00000407573 | TRUE | TRUE | FALSE |
| NPY1R | ENST00000296533 | TRUE | FALSE | FALSE |
| NPY2R | ENST00000329476 | TRUE | FALSE | FALSE |
| NPY4R | ENST00000374312 | TRUE | FALSE | FALSE |
| NR1D1 | ENST00000246672 | TRUE | FALSE | FALSE |
| NR1D2 | ENST00000312521 | TRUE | FALSE | FALSE |
| NR2E1 | ENST00000368986 | TRUE | FALSE | FALSE |
| NR2E3 | NA | FALSE | FALSE | FALSE |
| PAH | ENST00000553106 | TRUE | FALSE | FALSE |
| PBRM1 | ENST00000394830 | TRUE | TRUE | FALSE |
| PCSK2 | ENST00000262545 | TRUE | TRUE | FALSE |
| PDPK1 | ENST00000342085 | TRUE | TRUE | FALSE |
| PER1 | ENST00000317276 | TRUE | TRUE | FALSE |
| PER2 | ENST00000254657 | TRUE | TRUE | FALSE |
| PER3 | ENST00000361923 | TRUE | FALSE | FALSE |
| PHF10 | ENST00000339209 | TRUE | FALSE | FALSE |
| PIK3CA | ENST00000263967 | TRUE | TRUE | FALSE |
| PIK3CB | ENST00000477593 | TRUE | FALSE | FALSE |

| | | | | |
|---|---|---|---|---|
| PIK3CD | ENST00000377346 | TRUE | FALSE | FALSE |
| PIK3CG | ENST00000359195 | TRUE | FALSE | FALSE |
| PIK3R1 | ENST00000521381 | TRUE | TRUE | FALSE |
| PIK3R2 | ENST00000222254 | TRUE | FALSE | FALSE |
| PIK3R3 | ENST00000262741 | TRUE | FALSE | FALSE |
| PPARA | ENST00000396000 | TRUE | FALSE | FALSE |
| PPARD | ENST00000311565 | TRUE | FALSE | FALSE |
| PPARG | ENST00000287820 | TRUE | FALSE | FALSE |
| PPIA | ENST00000468812 | TRUE | TRUE | FALSE |
| PPIF | ENST00000225174 | TRUE | FALSE | FALSE |
| PPM1A | ENST00000325642 | TRUE | FALSE | FALSE |
| PPM1G | ENST00000344034 | TRUE | FALSE | FALSE |
| PRAF2 | ENST00000376390 | TRUE | FALSE | FALSE |
| PRKACA | ENST00000308677 | TRUE | TRUE | FALSE |
| PRKACB | ENST00000370685 | TRUE | TRUE | FALSE |
| PRKACG | ENST00000377276 | TRUE | FALSE | FALSE |
| PRKAR2A | ENST00000265563 | TRUE | TRUE | FALSE |
| PRKAR2B | ENST00000265717 | TRUE | TRUE | FALSE |
| PRKCE | ENST00000306156 | TRUE | FALSE | FALSE |
| PRKCH | ENST00000332981 | TRUE | FALSE | FALSE |
| PSMD1 | ENST00000308696 | TRUE | FALSE | FALSE |
| PTEN | ENST00000371953 | TRUE | TRUE | FALSE |
| RAB3A | ENST00000222256 | TRUE | FALSE | FALSE |
| RAB3B | ENST00000371655 | TRUE | FALSE | FALSE |
| RAB3C | ENST00000282878 | TRUE | TRUE | FALSE |
| RAB3D | ENST00000222120 | TRUE | FALSE | FALSE |
| RAC1 | ENST00000356142 | TRUE | TRUE | FALSE |
| RAC2 | ENST00000249071 | TRUE | TRUE | FALSE |
| RAC3 | ENST00000306897 | TRUE | FALSE | FALSE |
| REL | ENST00000295025 | TRUE | FALSE | FALSE |
| RELA | ENST00000406246 | TRUE | FALSE | FALSE |
| RELB | ENST00000221452 | TRUE | FALSE | FALSE |
| RFNG | ENST00000310496 | TRUE | TRUE | FALSE |
| RHBDL1 | ENST00000219551 | TRUE | FALSE | FALSE |
| RHBDL2 | ENST00000289248 | TRUE | FALSE | FALSE |
| RHBDL3 | ENST00000269051 | TRUE | FALSE | FALSE |
| RHEB | ENST00000262187 | TRUE | FALSE | FALSE |
| RHEBL1 | ENST00000301068 | TRUE | TRUE | FALSE |
| RHOA | ENST00000418115 | TRUE | TRUE | FALSE |
| RHOB | ENST00000272233 | TRUE | FALSE | FALSE |
| RHOC | ENST00000285735 | TRUE | FALSE | FALSE |
| RPS6KB1 | ENST00000225577 | TRUE | FALSE | FALSE |
| RPS6KB2 | ENST00000312629 | TRUE | FALSE | FALSE |
| RSU1 | ENST00000377921 | TRUE | FALSE | FALSE |
| SCAP | ENST00000265565 | TRUE | FALSE | FALSE |
| SDC1 | ENST00000381150 | TRUE | TRUE | FALSE |
| SDC2 | ENST00000302190 | TRUE | TRUE | FALSE |
| SDC3 | ENST00000339394 | TRUE | FALSE | FALSE |
| SDC4 | ENST00000372733 | TRUE | TRUE | FALSE |
| SGK1 | ENST00000367858 | TRUE | TRUE | FALSE |
| SGK2 | ENST00000341458 | TRUE | FALSE | FALSE |
| SIRT1 | ENST00000212015 | TRUE | FALSE | FALSE |
| SIRT3 | ENST00000382743 | TRUE | FALSE | FALSE |
| SLC18A1 | ENST00000440926 | TRUE | FALSE | FALSE |
| SLC18A2 | ENST00000298472 | TRUE | TRUE | FALSE |
| SLC18A3 | ENST00000374115 | TRUE | FALSE | FALSE |
| SLC32A1 | ENST00000217420 | TRUE | TRUE | FALSE |
| SLC6A2 | ENST00000219833 | TRUE | FALSE | FALSE |
| SLC6A3 | ENST00000270349 | TRUE | FALSE | FALSE |
| SLC9A3R1 | ENST00000262613 | TRUE | TRUE | FALSE |
| SLC9A3R2 | ENST00000424542 | TRUE | TRUE | FALSE |
| SMARCA2 | ENST00000382203 | TRUE | TRUE | FALSE |

| | | | | |
|---|---|---|---|---|
| SMARCA4 | ENST00000429416 | TRUE | TRUE | FALSE |
| SMARCC1 | ENST00000254480 | TRUE | TRUE | FALSE |
| SMARCC2 | ENST00000267064 | TRUE | FALSE | FALSE |
| SMARCD1 | ENST00000394963 | TRUE | TRUE | FALSE |
| SMARCD2 | ENST00000448276 | TRUE | TRUE | FALSE |
| SMARCD3 | ENST00000262188 | TRUE | TRUE | FALSE |
| SMARCE1 | ENST00000348513 | TRUE | FALSE | FALSE |
| SREBF1 | ENST00000355815 | TRUE | TRUE | FALSE |
| SREBF2 | ENST00000361204 | TRUE | TRUE | FALSE |
| STX1A | ENST00000222812 | TRUE | TRUE | FALSE |
| STX1B | ENST00000215095 | TRUE | TRUE | FALSE |
| STX2 | ENST00000392373 | TRUE | TRUE | FALSE |
| STX3 | ENST00000337979 | TRUE | TRUE | FALSE |
| STX4 | ENST00000313843 | TRUE | FALSE | FALSE |
| STXBP1 | ENST00000373302 | TRUE | TRUE | FALSE |
| STXBP2 | ENST00000221283 | TRUE | FALSE | FALSE |
| STXBP3 | ENST00000370008 | TRUE | FALSE | FALSE |
| SYN1 | ENST00000295987 | TRUE | FALSE | FALSE |
| SYN2 | NA | FALSE | TRUE | FALSE |
| SYN3 | ENST00000358763 | TRUE | FALSE | FALSE |
| TAF4 | ENST00000252996 | TRUE | FALSE | FALSE |
| TAF4B | ENST00000269142 | TRUE | FALSE | FALSE |
| TAOK1 | ENST00000261716 | TRUE | TRUE | FALSE |
| TAOK2 | ENST00000308893 | TRUE | FALSE | FALSE |
| TAOK3 | ENST00000392533 | TRUE | FALSE | FALSE |
| TBX20 | ENST00000408931 | TRUE | FALSE | FALSE |
| TH | ENST00000381178 | TRUE | TRUE | FALSE |
| TIMELESS | ENST00000553532 | TRUE | FALSE | FALSE |
| TLR1 | ENST00000308979 | TRUE | FALSE | FALSE |
| TLR10 | ENST00000308973 | TRUE | FALSE | FALSE |
| TLR2 | ENST00000260010 | TRUE | FALSE | FALSE |
| TLR3 | ENST00000296795 | TRUE | FALSE | FALSE |
| TLR4 | ENST00000355622 | TRUE | FALSE | FALSE |
| TLR5 | ENST00000540964 | TRUE | FALSE | FALSE |
| TLR6 | ENST00000436693 | TRUE | FALSE | FALSE |
| TLR7 | ENST00000380659 | TRUE | FALSE | FALSE |
| TLR8 | ENST00000218032 | TRUE | FALSE | FALSE |
| TLR9 | NA | FALSE | FALSE | FALSE |
| TPH1 | ENST00000250018 | TRUE | FALSE | FALSE |
| TPH2 | ENST00000333850 | TRUE | TRUE | FALSE |
| TRPV1 | ENST00000571088 | TRUE | FALSE | FALSE |
| TRPV2 | ENST00000338560 | TRUE | FALSE | FALSE |
| TRPV3 | ENST00000301365 | TRUE | FALSE | FALSE |
| TRPV4 | ENST00000418703 | TRUE | FALSE | FALSE |
| TRPV5 | ENST00000265310 | TRUE | FALSE | FALSE |
| TRPV6 | ENST00000359396 | TRUE | FALSE | FALSE |
| TSPO | ENST00000329563 | TRUE | FALSE | FALSE |
| TSPO2 | ENST00000373161 | TRUE | FALSE | FALSE |
| UGDH | ENST00000316423 | TRUE | FALSE | FALSE |
| UNC13A | ENST00000519716 | TRUE | FALSE | FALSE |
| UNC13B | ENST00000378495 | TRUE | FALSE | FALSE |
| UNC13C | ENST00000260323 | TRUE | FALSE | FALSE |
| UNC79 | ENST00000256339 | TRUE | FALSE | FALSE |
| UNC80 | ENST00000439458 | TRUE | FALSE | FALSE |
| XRCC5 | ENST00000392133 | TRUE | FALSE | FALSE |

**Supplemental Section S1:**

Simulation Details:

The simulation LOF, SYN, and MIS variant count data was generated using the corrvar() function from the *SimCorMix* R package according to the following distributions:

$LOF\ count \sim Neg\ Binom\ (size\ =\ 1.25, prob\ =\ 0.881)$,

$SYN\ count \sim Neg\ Binom\ (size\ =\ 2.5, prob\ =\ 0.638)$, and

$MIS\ count \sim Zero - Inflated\ Neg\ Binom\ (size\ =\ 8, prob\ =\ 0.73, prob\ str\ zero\ =\ 0.2)$,

using observed correlation matrix [insert matrix in Word].

$$\begin{bmatrix} 1 & 0.166 & 0.257 \\ 0.166 & 1 & 0.688 \\ 0.257 & 0.688 & 1 \end{bmatrix}$$

The GOI outcome was generated such that:

$$P(GOI)\ =\ \frac{1}{(1 + exp(-1 * (Int + LOF + SYN + MIS))}$$

If $P(GOI)\ <\ 0.5, Control, else, GOI.$

The parameters used to generate the random data for all runs are shown in **Supplemental Table 3**. To assess the performance of the approach, we compared the parameter estimates from each model to these true parameter values used to generate the response. In total 18 separate simulation scenarios were designed representing 3 different beta parameter vectors and a range of standard deviations used in the random error distribution. Each scenario was iterated across 1000 randomly generated datasets. The performance of each model

**Supplemental Table S3:** Parameters used to generate random data for the simulations.

| Run | Min GOI | Max GOI | Med GOI | LOF | SYN | MIS | Err sd |
|---|---|---|---|---|---|---|---|
| 1a | 53 | 113 | 83 | 0.75 | 0.1 | 0.05 | 0.75 |
| 1b | 86 | 158 | 123 | 0.75 | 0.1 | 0.05 | 0.85 |
| 1c | 139 | 229 | 183 | 0.75 | 0.1 | 0.05 | 0.95 |
| 2a | 24 | 61 | 39 | 1.25 | 0.2 | 0.1 | 0.65 |
| 2b | 26 | 68 | 44 | 1.25 | 0.2 | 0.1 | 0.75 |
| 2c | 31 | 74 | 52 | 1.25 | 0.2 | 0.1 | 0.85 |
| 3a | 59 | 122 | 88 | 1.75 | 0.3 | 0.2 | 1.25 |
| 3b | 66 | 131 | 99 | 1.75 | 0.3 | 0.2 | 1.35 |
| 3c | 80 | 146 | 110 | 1.75 | 0.3 | 0.2 | 1.45 |

Legend:
Min/Max/Med GOI: minimum/maximum/median number of GOIs in the random datasets generated under the given run parameters
LOF: True LOF parameter value used to generate the random response
SYN: True SYN parameter value used to generate the random response
MIS: True MIS parameter value used to generate the random response
Err sd: Standard deviation used in the distribution of the random error

**Supplemental Table S4:** Parameter estimates from simulations.

| Run | min COI | max COI | med COI | LOFest | SYNest | MISest | LOFz | SYNz | MISz |
|---|---|---|---|---|---|---|---|---|---|
| 1a | 731 | 1754 | 1173 | 2.598 (0.0085) | 0.353 (0.0042) | 0.184 (0.0034) | 10.193 (0.0222) | 2.923 (0.0327) | 1.886 (0.0341) |
| 1b | 1180 | 2253 | 1683.5 | 2.199 (0.0058) | 0.295 (0.003) | 0.162 (0.0024) | 12.382 (0.022) | 3.333 (0.033) | 2.286 (0.033) |
| 1c | 1745 | 3115 | 2406 | 1.884 (0.0043) | 0.252 (0.0023) | 0.141 (0.0018) | 14.181 (0.0239) | 3.727 (0.0334) | 2.642 (0.0331) |
| 2a | 452 | 1317 | 880 | 3.477 (0.019) | 0.569 (0.0067) | 0.285 (0.0056) | 7.106 (0.0269) | 2.984 (0.0275) | 1.815 (0.0318) |
| 2b | 588 | 1549 | 1051 | 3.065 (0.0121) | 0.502 (0.0051) | 0.254 (0.0043) | 8.575 (0.0235) | 3.337 (0.0294) | 2.061 (0.0322) |
| 2c | 774 | 1746 | 1251.5 | 2.761 (0.0094) | 0.447 (0.0041) | 0.232 (0.0033) | 10.039 (0.0219) | 3.652 (0.0301) | 2.317 (0.0318) |
| 3a | 753 | 1770 | 1239 | 3.413 (0.0139) | 0.589 (0.0051) | 0.399 (0.0045) | 8.348 (0.0245) | 3.84 (0.0263) | 3.087 (0.0289) |
| 3b | 870 | 1925 | 1376.5 | 3.15 (0.0113) | 0.54 (0.0043) | 0.371 (0.0038) | 9.365 (0.0232) | 4.062 (0.0273) | 3.326 (0.0289) |
| 3c | 990 | 2065 | 1522.5 | 2.929 (0.0095) | 0.501 (0.0037) | 0.344 (0.0031) | 10.389 (0.0215) | 4.297 (0.0281) | 3.529 (0.0285) |

Legend:
min/max/medCOI: minimum, maximum, and median number of genes in the clusters of interest
LOF/SYN/MISest: mean parameters estimate and standard error of those means for the LOF, SYN, and MIS parameters
LOF/SYN/MISz: mean z-score and standard error of those scores for the estimated effect of the LOF, SYN, and MIS parameters
